# PERSISTENT LOW-LEVEL INFECTIONS OF ELEPHANT ENDOTHELIOTROPIC HERPESVIRUSES AND ELEPHANT GAMMAHERPESVIRUSES DETECTED IN SKIN NODULES AND SALIVA FROM WILD AND ZOO AFRICAN ELEPHANTS

**DOI:** 10.64898/2026.04.14.718412

**Authors:** Virginia Riddle Pearson, Gary S. Hayward

## Abstract

The goal of this study was to determine which species of Elephant Endotheliotropic Herpesviruses (EEHV) and Elephant Gammaherpesviruses (EGHV) are endemic in wild African elephants, *Loxodonta*. We collected skin nodule biopsies, saliva and tissues from 43 wild *L. africana* (African bush elephant) in Botswana, Kenya, South Africa and Zimbabwe; and saliva from 25 wild *L. cyclotis* (African forest elephant) in Gabon. We also collected saliva over seven years from 7 wild-born *L. africana* at Six Flags Safari Park, USA, and saliva, blood and tissues from an additional 200 *L. africana* and *Elephas maximus* in USA zoos. Conventional polymerase chain reaction and Sanger sequencing of purified DNA from these samples yielded thousands of unambiguous positive genetic matches to known EEHV2, EEHV3A, EEHV3B, EEHV6, EEHV7A, EGHV2, EGHV3B, EGHV4B, EGHV5B, and discovered new species EEHV3C-H, EEHV7B and EGHV1B in African elephants, and EGHV5A in an Asian elephant. Our extensive library of EEHV and EGHV sequences from wild and zoo elephants provide a significant resource to the elephant virologists and conservationists.

## INTRODUCTION

The last of the ancient Order Proboscidea: Genus *Loxodonta*, including *L. africana,* (African bush elephant) and *L. cyclotis* (African forest elephant), and Genus *Elephas,* including *Elephas maximus* (Asian elephants) are listed by the IUCN as highly or critically endangered. [1,2,]. Habitat fragmentation, increasing human-elephant conflict, and poaching of elephants for ivory, skin and internal organs have decimated entire populations in all traditional range countries. [3–10]. *Loxodonta* and *Elephas* are further threatened by lethal Elephant Hemorrhagic Disease (EHD) caused by Elephant Endotheliotropic Herpesviruses (EEHV). Hundreds of confirmed cases of lethal or severe EHD attributed to EEHV1A, EEHV1B, EEHV4 and EEHV5 have been documented in wild and zoo *E. maximus*, *E.m borneensis* and *E.m sumantranus.* [11–34]. Nearly two dozen cases of lethal or severe EHD attributed to EEHV2, EEHV3A, EEHV3B, and EEHV6, but not EEHV7A or EEHV7B, have been documented in *L. africana* in zoos worldwide. However, no EHD cases have been reported to date in range countries. [35–42]. As the numbers of free-ranging *Loxodonta* continue to decline precipitously, and wild herds become genetically isolated, lethal EHD may become a much more significant threat. [43]. Previously, we reported finding sub-clinical evidence of EEHV 2, EEHV3A, EEHV6, and EEHV7A in lung biopsies from a poached wild *L. africana* in Kenya (GenBank accession #KT832496-KT832512). [44]. **Fig 1**.

**Figure 1.**
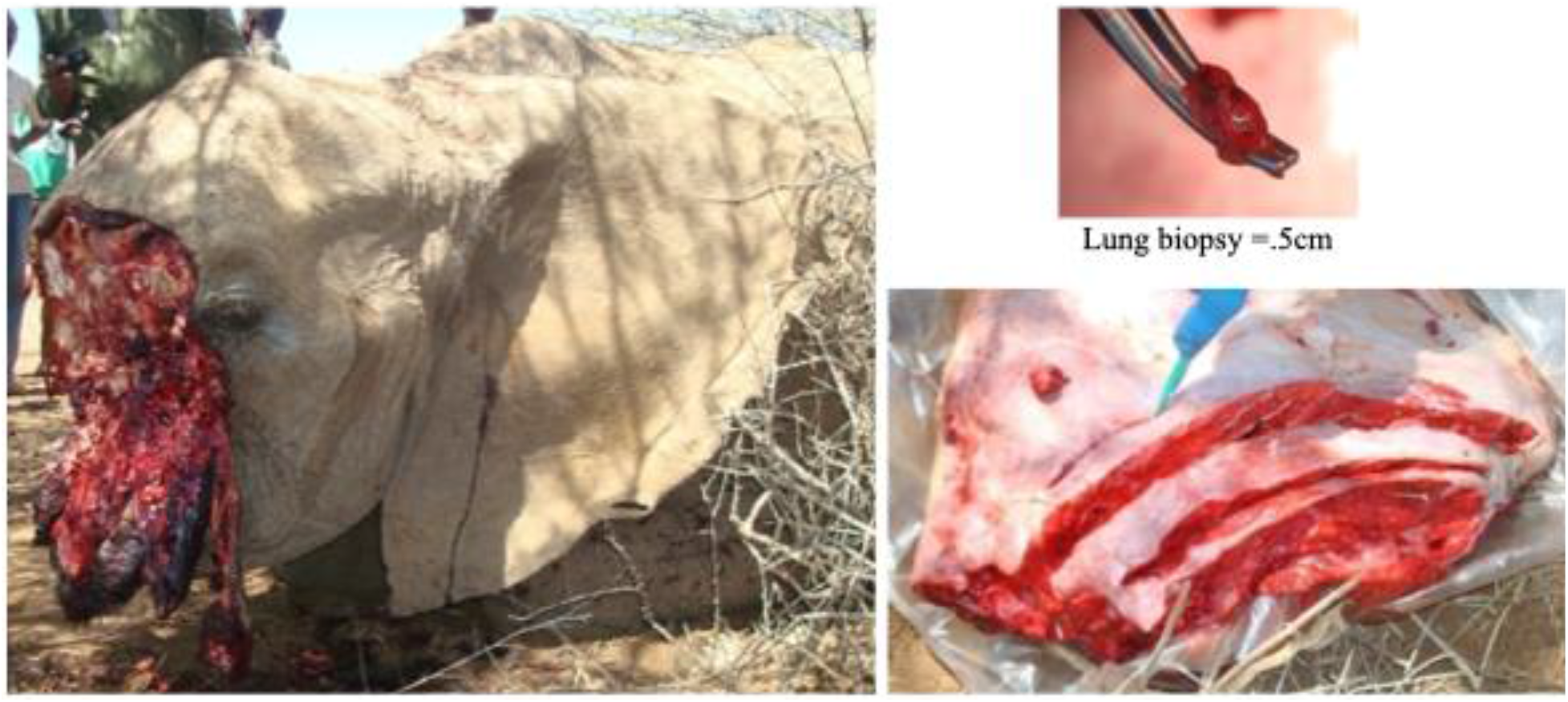
left, wild *L. africana* adult female, just outside Samburu National Reserve, Kenya; bottom right, lung collected at field necropsy by Pearson; top right, .5 cm lung biopsies were immersed in Qiagen RNAlater or Pax Gene Tissue Container, credit Pearson, 2011.

The goal of this study was to determine which species of EEHV and EGHV are endemic in wild *Loxodonta*. Descriptions of viral nuclear inclusion bodies morphologically consistent with herpesviruses in raised cutaneous fibropapillomas (skin nodules) had been reported in a group of ninety-nine *L. africana* imported to USA from Zimbabwe in 1982-84 [45], as were findings of white lymphoid nodules containing Cowdry Type A intranuclear inclusions pathognomic of herpesvirus disease in lungs of culled *L. africana* in South Africa. [46]. We hypothesized that we might find evidence of since-characterized EEHV and EGHV in similar skin nodules and lungs in living wild *Loxodonta* to discover unique genetic polymorphisms that could help understand the origins of EHD. **Fig 2**.

**Figure 2.**
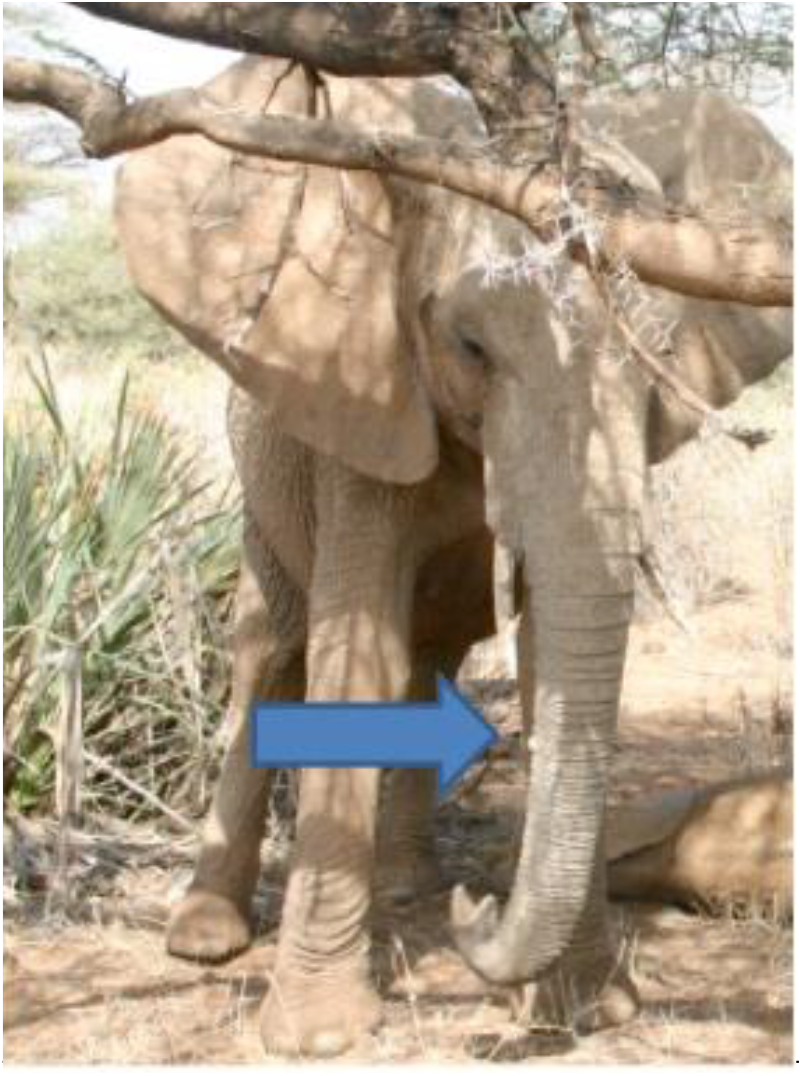
Skin nodules observed on wild juvenile *L. africana,* Samburu National Reserve, Kenya, credit Pearson, 2009

In 2011 and 2013, in collaboration with Save The Elephants and Kenya Wildlife Service Veterinary and Capture Service in Kenya; Elephants Without Borders in Botswana and Zimbabwe; and Elephants Alive in South Africa, Pearson collected biopsies of skin nodules and saliva from three immobilized wild young *L. africana* calves (**Fig 3, 4**) and two subadults **(Fig 5**) in Kenya; and two immobilized wild young *L. africana* calves in Botswana (**Fig 6,7,8),** and saliva and non-nodular skin warts **(Fig. 9)** from twenty-six wild or semi-tame *L.africana* who had been immobilized for emergency treatment of gunshot wounds, GPS collaring operations or who were semi-habituated to humans. [47–50]. Because elephants can be immobilized for only a short time, we were unable to collect skin adjacent to the nodules for investigation of EEHV and EGHV cellular tropism. [51]. Pearson was also given skin nodule biopsies and/or saliva collected previously from six wild *L.africana* by South Africa National Parks Veterinary Unit. Samples were hand-carried or shipped to USA under USDA and CITES Permits. In 2017, Agence Nationale des Parcs Nationaux and CENEREST (National Centre for Scientific and Technological Research), Gabon, exported to us saliva swab samples from 25 *L. cyclotis* that had been immobilized for GPS collaring purposes.

**Figure 3.**
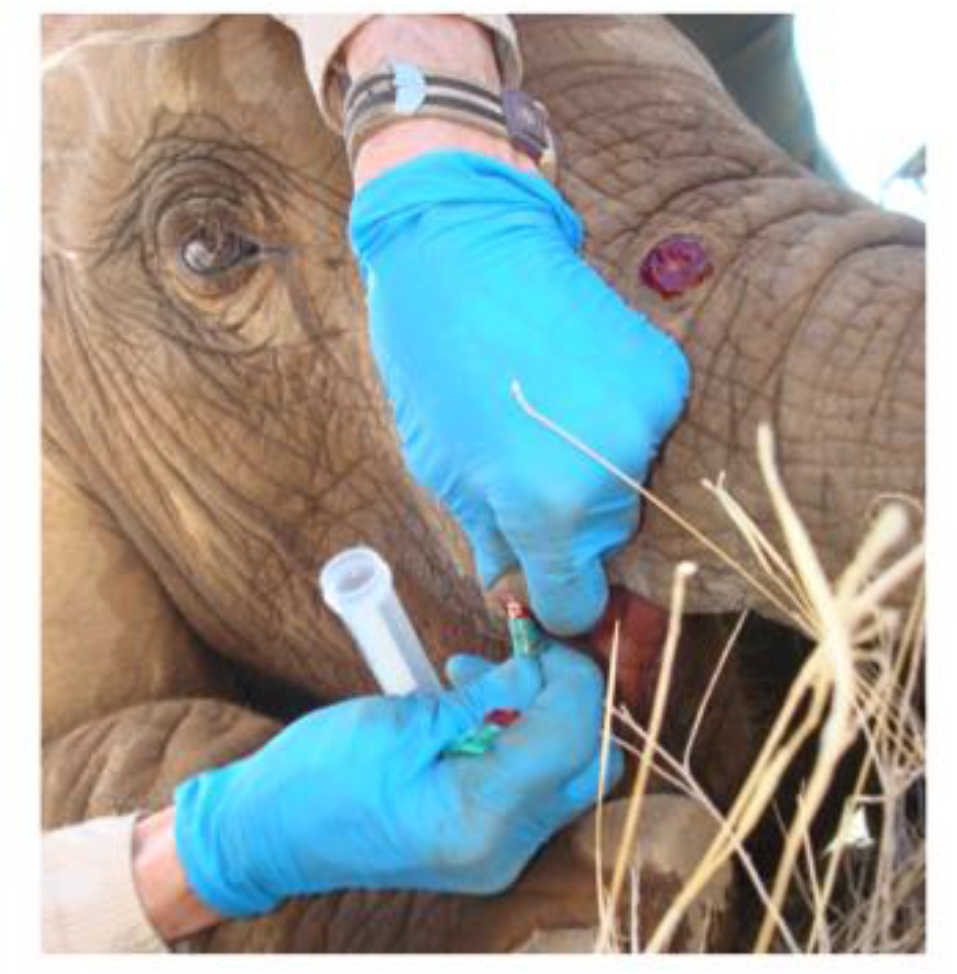
Skin nodule on immobilized wild *L. africana* female calf cGM, Samburu National Reserve, Kenya, credit Pearson, 2011.

**Fig 4.**
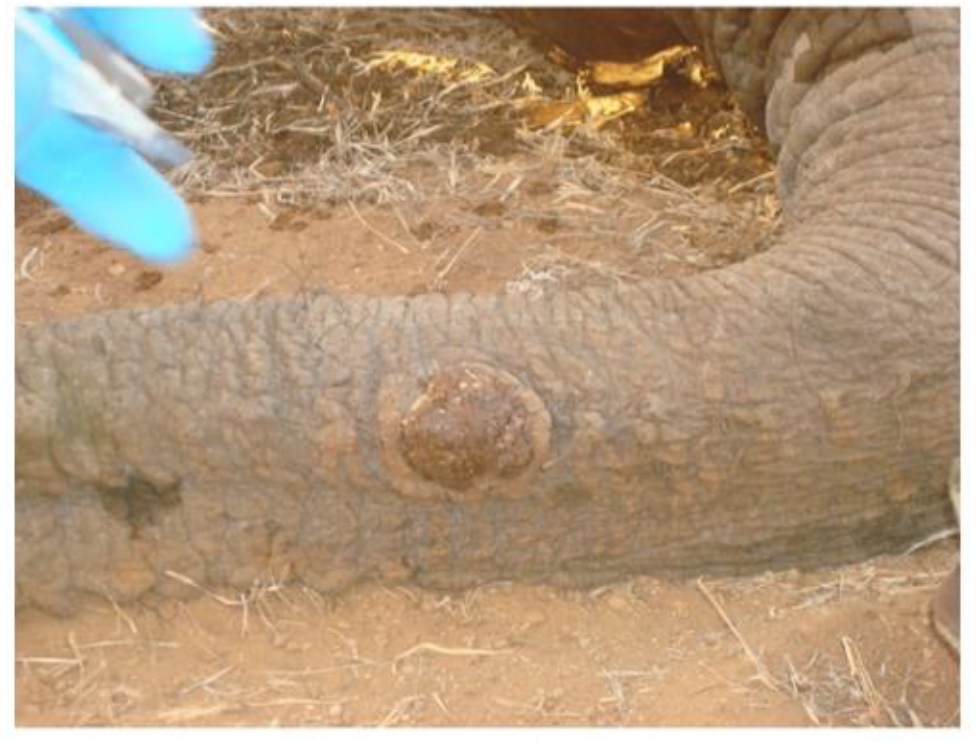
Skin nodule on immobilized wild *L. africana* female calf cMT, Samburu National Reserve, Kenya, credit Pearson, 2011.

**Fig 5.**
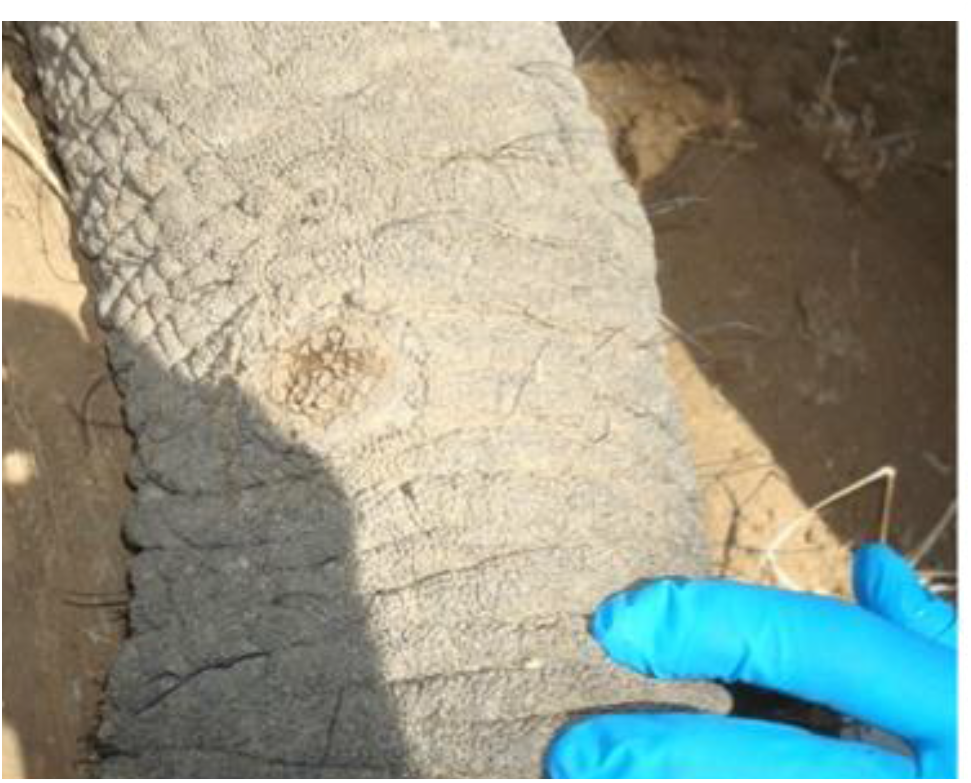
Crusted over skin nodule on immobilized wild *L. africana* juvenile female HIMA, Samburu National Reserve, Kenya, credit Pearson, 2011.

**Figure 6.**
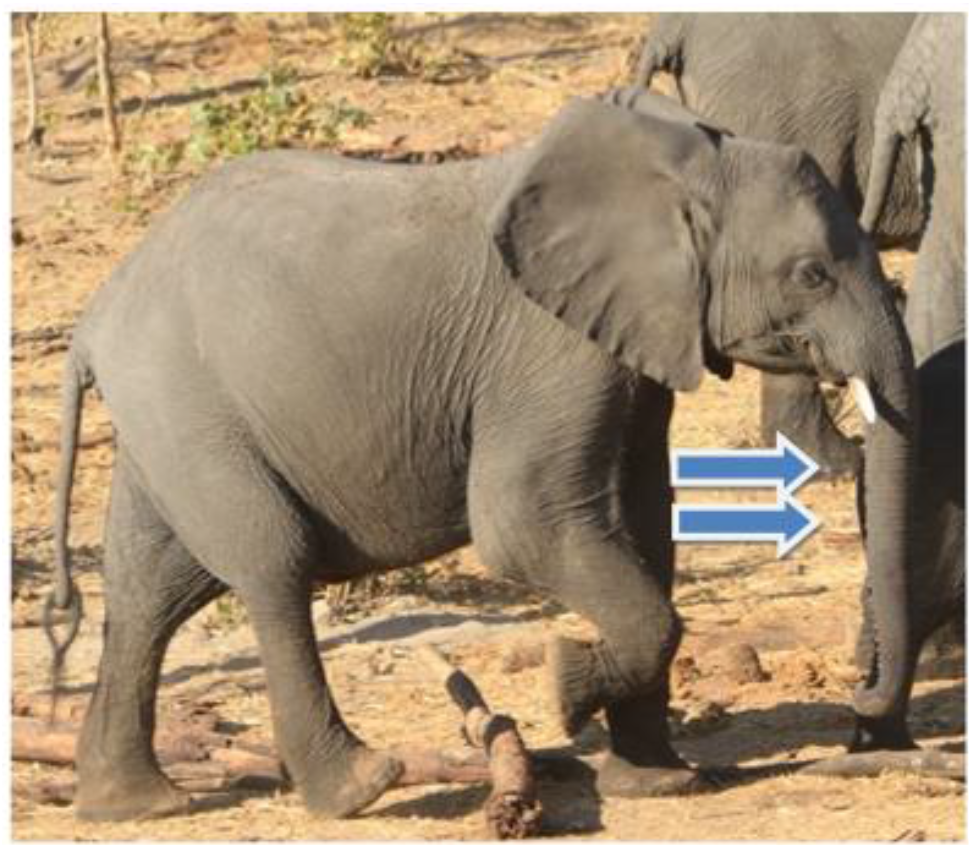
Four nodules on trunk underside of wild juvenile male *L.africana*, designated BW1M Nod1 to Nod 4. Chobe National Park, Kasane, Botswana, credit Pearson, 2013.

**Figure 7.**
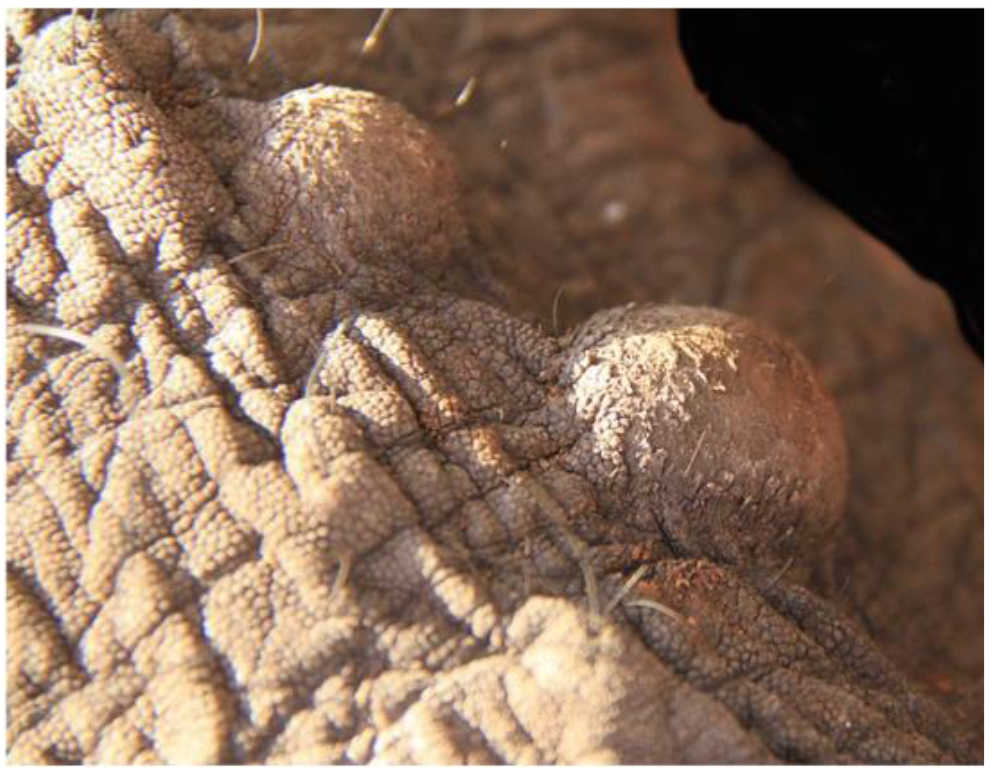
Close-up of skin nodules on wild *L. africana* young juvenile male BW1M Nod1, credit Pearson, 2013

**Figure 8.**
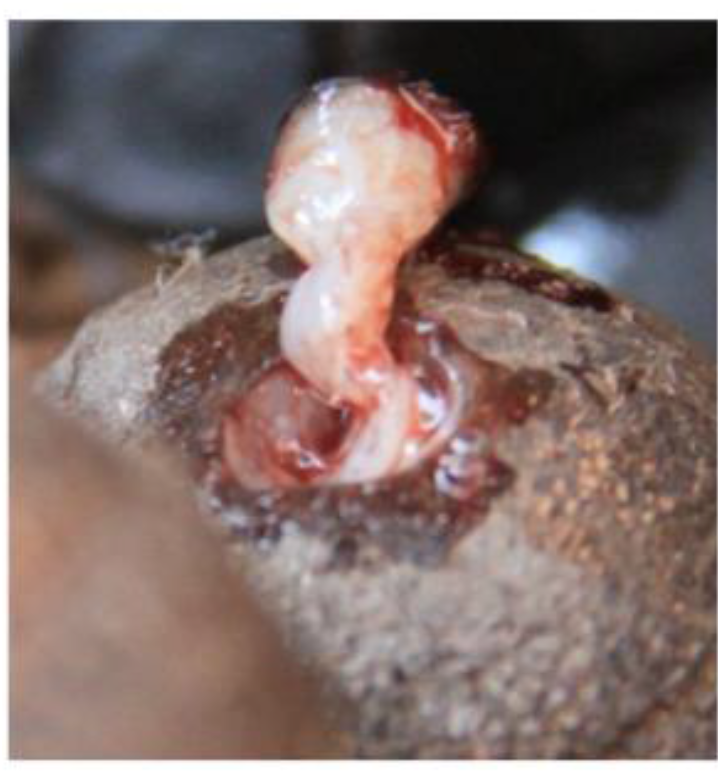
Biopsy (6mm) of skin nodule BW1M NOD1 (and NODS 2,3,4) were collected by Pearson and immersed in RNALater in Chobe National Park, Botswana, credit Pearson, 2013.

**Figure 9.**
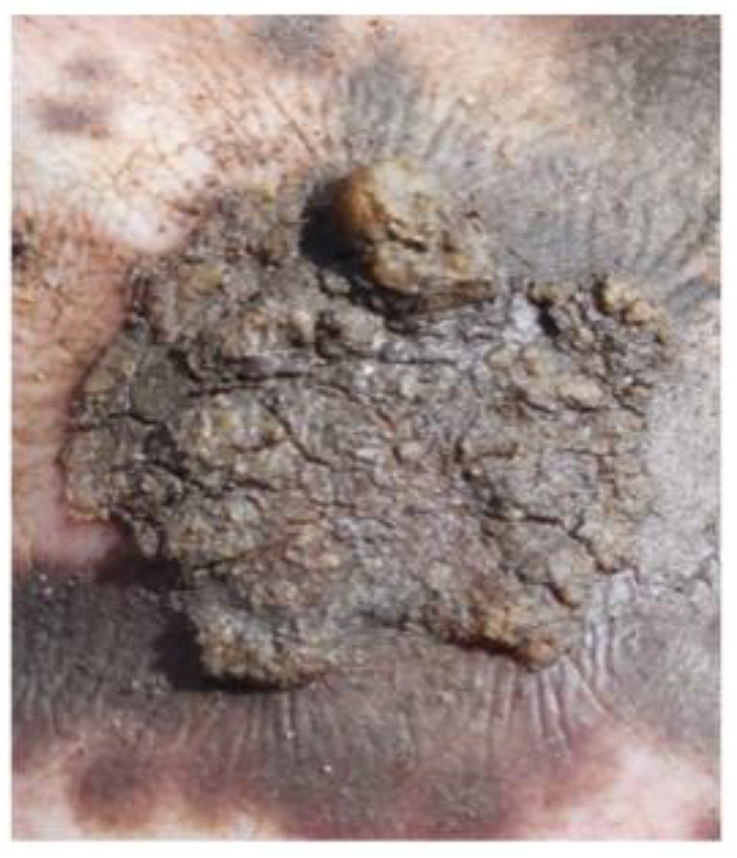
Punch biopsies (5 mm) of ear warts on immobilized wild *L. africana* adult bull RSA Wild Spirit, Balule, South Africa, were preserved in Qiagen RNAlater, credit Pearson, 2013.

Concurrently, we investigated the frequency and types of EEHV and EGHV shed in saliva over seven years (2012-2019) from a herd of seven *L. africana* at Six Flags Safari Park, USA. In 2012, bi-weekly saliva swabs were collected from 2 adult female *L. africana* (SFL and SFJ) and periodically from the entire herd through 2019 for a total of 180 saliva samples. SFL, the herd matriarch, was imported from Uganda in 1971-72 and had lived at the park ever since.

SFJ was a survivor of the ninety-nine *L. africana* imported from Zimbabwe in 1982-84 [45] and had lived at several other USA zoos before coming to Six Flags in 2010. The other five herd mates, imported from Uganda with the matriarch, had never left the safari park. It is important to note that all Six Flags *L. africana* were asymptomatic for EHD at the onset and throughout this study. Our novel technique of collecting saliva by extra-long protological swabs was effective to sample anaesthetized wild elephants or elephants such as these who were not accustomed to routine trunk wash or blood collection protocols used in many USA zoos to detect systemic EHD viremia. [52–57]. One caveat of this seven-year saliva study is that we have no saliva samples from any of the herd members prior to the introduction of the Zimbabwe-born SFJ to address the question of viral cross-transmission within the herd, **Fig 10**.

**Fig 10.**
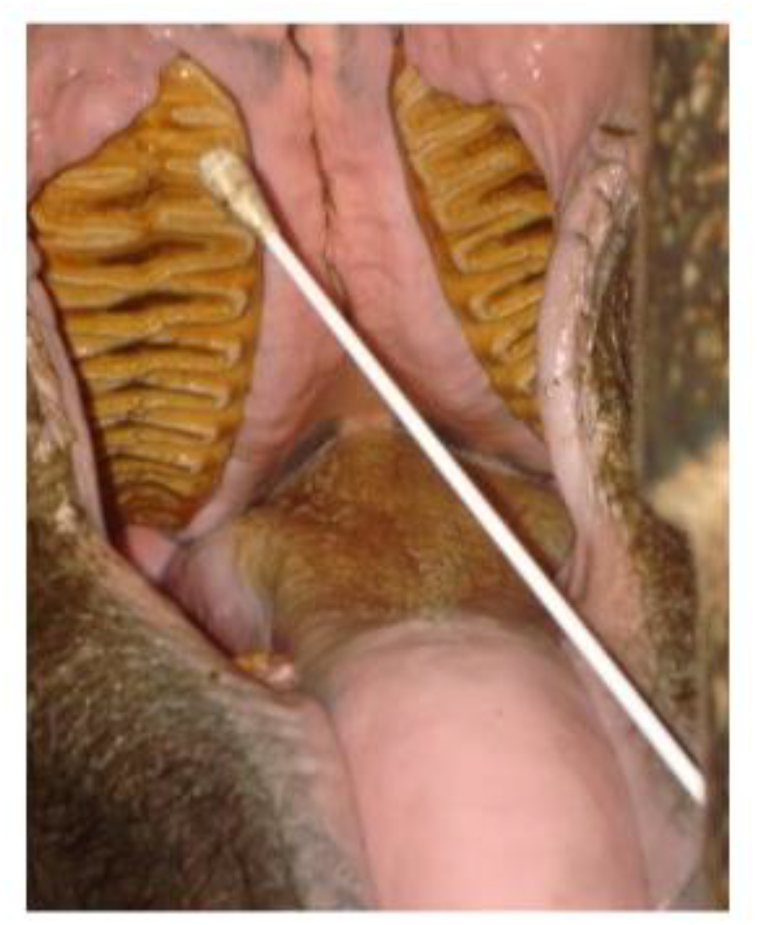
*L.africana* adult female SFJ, Six Flags Safari Park, USA, credit Pearson, 2012.

Although not as readily subject to quantitative analysis (because of considerably more variability in individual elephant and handler collection techniques and recovered saliva volumes), in a similar study, this saliva swab plus Sanger PCR subtype DNA sequencing approach proved to closely parallel the results obtained in trunk wash procedure for detecting EEHV3B in a young USA zoo *L. africana* recuperating from EHD. [35]. The relative abundance of viral DNA obtained by the two approaches was not addressed but scoring of positive results seemed to be at least equally sensitive with both methods being able to detect the continued presence of viral DNA out to about eight months. The results also demonstrated that no significant change occurred in the EEHV3B DNA sequence obtained at each of three PCR loci over this post-viremia monitoring period. Further investigation of saliva using quantitative polymerase chain reaction (qPCR) is needed to evaluate saliva as a useful diagnostic and prognostic tool for early indication of EHD as compared to blood and trunk wash. [58,59].

## METHODS AND MATERIALS

### PERMITS AND APPROVALS

Field sample collections were conducted under Elephants Without Borders Botswana Research Permit/ #WT8/36/4XV(41 (GPS-coordinates Chobe National Park, Kasane, Botswana, 17°47’42.83"S, 25°10’16.067"E); Kenya Research Permit #NCST/RRI/12/1MAS/9 Permittee Virginia R. Pearson; Save The Elephants Research Permit with Kenya Service Veterinary Capture and Services Department (GPS-coordinates Samburu National Reserve, Samburu County, Kenya, 0°37’5”N, 37°31’48”E and Keekorok, Masai Mara National Reserve, Narok, Kenya, 1°35’ 9.00"S, +35° 15’ 6.00"E); and Elephants Alive with South African National Parks Veterinary Wildlife Services, (GPS-coordinates Timbavati Reserve, Mpumalanga Province, South Africa, 24°20’07S,31°20’38”E, and Crocodile Bridge, Kruger National Park, South Africa, 25°21’0”S,31°53’32”E); and CENEREST(National Centre for Scientific and Technological Research) and Agence Nationale des Parcs Nationaux Gabon, (GPS-coordinates Minkebe NP 1°40’47”N,12°45’34”E; Mwagna NP 0°36’N,12°42’E; Ivindo NP 0.088°N, 12.63°E; Pongara NP 0°07’N,9°38’E; Loango NP 2°10’S,9°34’E; Moukalaba Doudou NP 2°26’S, 10°25’E). Samples were imported into USA under United States Veterinary Permit for Importation and Transportation of Controlled Materials and Organisms and Vectors #11798, Permittee Virginia R. Pearson; San Diego Zoo Global CITES/ USFWS Import Permit # 13-US727416/9 and #10-US727416/9; International Elephant Foundation CITES/USFWS Import Permit #17US09806C/9; Botswana CITES Export Permit #0202118; Gabon CITES Export Permit #0785; Kenya CITES Export Permit # 008624/ OR82872; and South Africa CITES Export Permit #1087577. Protocols for elephant sample collection and laboratory analyses were approved by San Diego Zoo Global Institutional Animal Care and Use Committee IACUC #11– 002; Princeton University IACUC Enquist Laboratory Protocol #1819; and Fox Chase Cancer Center Rall Laboratory federal regulations governing research involving material of animal origin (elephant). Samples from within the United States were collected according to participating zoos’ Animal Welfare and Research Committees and Association of Zoos and Aquariums Elephant Research and Necropsy Protocol.

### SKIN NODULE BIOPSY AND SALIVA COLLECTION

Elephants were immobilized by qualified, experienced field veterinarians using etorphine hydrochloride (M99). 5 mm punch biopsies were excised from trunk nodules and immersed in 2.5 ml RNALater Reagent (Qiagen). Incisions were treated with iodine and oxytetracycline. Diprenorphrine hydrochloride or naltrexone reversed sedation within 90 sec. No injuries to elephants or personnel occurred during these field operations. Saliva samples from immobilized, or semi-captive wild or captive elephants in Africa and America were collected using 16 in polyester swabs (Puritan Medical Corporation) until saturated then immersed in 2 ml RNAprotect Cell Reagent or RNALater (Qiagen) or pressed onto Gentegra DNA Collection GenSaver Cards.

### DNA EXTRACTION

Pearson worked with these samples first at Enquist Laboratory, Department of Molecular Biology, Princeton University, Princeton, NJ, USA, and subsequently at Rall Laboratory, Fox Chase Cancer Center, Philadelphia PA USA. Neither lab had previously ever worked with elephant herpesviruses. DNA was extracted from skin nodule biopsies, saliva, and tissues using Qiagen DNeasy Blood and Tissue Kit according to the manufacturer’s instructions with the modification of extended overnight incubation at 56’C, followed by 10 min incubation at 95°C. Qiagen Repli-G was used to amplify extracted DNA in cases when three-round nested cPCR was necessary to detect expected very low levels of viral genomes.

### CONVENTIONAL POLYMERASE CHAIN REACTION (cPCR): SEE SUPPLEMENTAL INFORMATION (SI) FOR PRIMER LIST

DNA was amplified by cPCR using three-round nested primer sets targeting EEHV genes loci including U38(POL), U39(gB), U47(gO), U66(TERex3), U77(HEL), U48.5(TK), U51(vGPCR1, U71/gM, U73(OBP), U77(HEL), U81(UDG), U82(gL), E24(vOX2B1), E4(vGCNT1), E37(ORF-0ex3), E9A(vOGT), E16D(vECTL), E54(vOX2-1), U14, U42(MTA) and U43(PRI) and primers designed specifically to recognize EGHV genes loci. Many other primer combinations were used to distinguish unique polymorphisms of subtypes EEHV3A-EEHV3H, including chimeric domaines CD-I U39(gB), CD-II U47(gO), U48.5(TK), U48(gH), CD-III U81(UDG), U82(gL) E37(ORF-Oex3) CD IV E9A(vOCT) CD-V E16D(vECTL), among others. For each 25 µl cPCR reaction, we used 2 µl elephant DNA, 12.5 µl GoTaq G2 Hot Start Green Master Mix (Promega), 8 µl sterile nuclease-free H2O, and 1.25 µl each forward and reverse primers. Reactions were run on an Eppendorf MasterCycler or Finnzymes PIKO Thermocycler programmed at 95°C for 2 min; 45 cycles of 95°C for 40 sec, 50°C for 45 sec; 73°C for 60 sec; final extension 78°C for 7 min. 25 µl of each PCR reaction was run on 2% agarose electrophoresis gels with ethidium bromide or GelRed or GelGreen nucleic acid stain (Phenix Research Products). Bright bands of expected, and often unexpected size, were purified and sent for sequencing.

### SANGER SEQUENCING

Primer extension sequencing was performed by Genewiz, Inc (South Plainfield, NJ) using Applied Biosystems BigDye version 3.1. The reactions were then run on Applied Biosystem 3730xl DNA Analyzer.

### CONTAMINATION CONTROLS

Strict contamination protocols were followed including: 1) one tube containing sterile H20 or PBS in every cPCR run); 2) frequent cPCR testing of all reagents and routine replacement of reagents; 3) retesting in a remote laboratory of any cPCR reactions showing suspicious positives space using new reagents; 4) duplicate DNA extractions were performed in remote lab spaces if enough original sample remained. For independent DNA extraction, cPCR and Sanger sequence verification, a subset of skin nodule biopsies (Fig 12) was sent to the Hayward Laboratory, Johns Hopkins University School of Medicine. All EEHV and EGHV sequences generated at Princeton University and Fox Chase Cancer Center Laboratories were evaluated, verified and/or duplicated at Hayward Laboratory, **Fig 11**.

**Figure 11.**
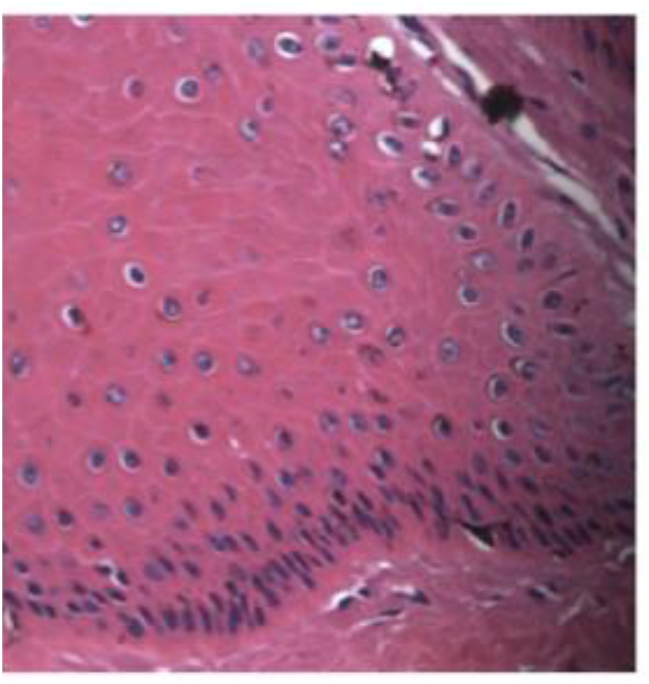
Biopsy of BW1M NOD1 immersed at collection in Qiagen PAXgene Tissue Container, was fixed in formalin-fixed paraffin preserved (FFPE) block, cut at 5 microns and stained with hematoxylin and eosin. Histological tissue slide shows multiple keratocyte nuclei that are expanded by a dark basophilic material (intranuclear inclusions indicative of viral infection) distorting the nuclei into an oval to rounded appearance surrounded by a 1-2 um clear halo, mag 45x, credit SY Long, Hayward Laboratory, Johns Hopkins School of Medicine.

**Fig 12.**
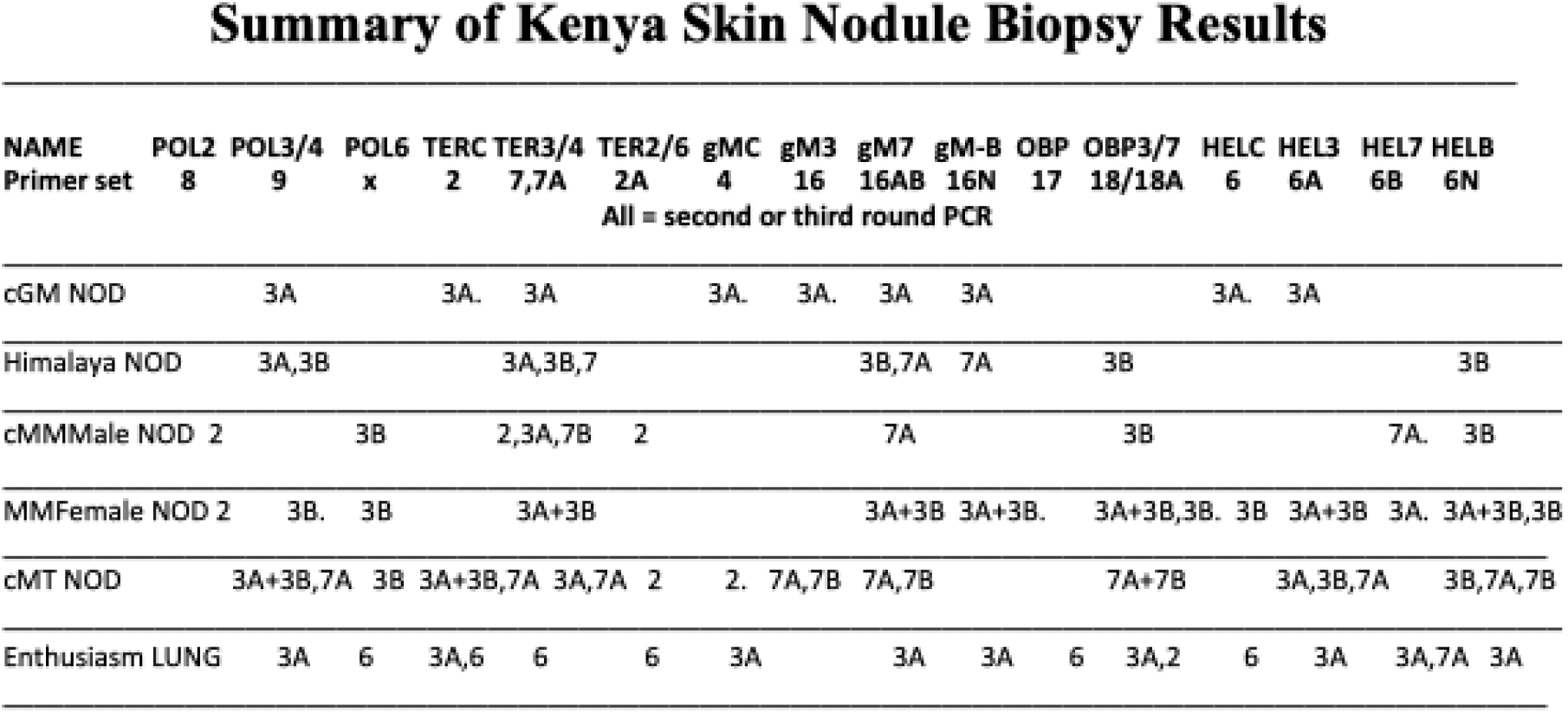
Summary of Kenya Skin Nodule Biopsy Results.

**Fig 13.**
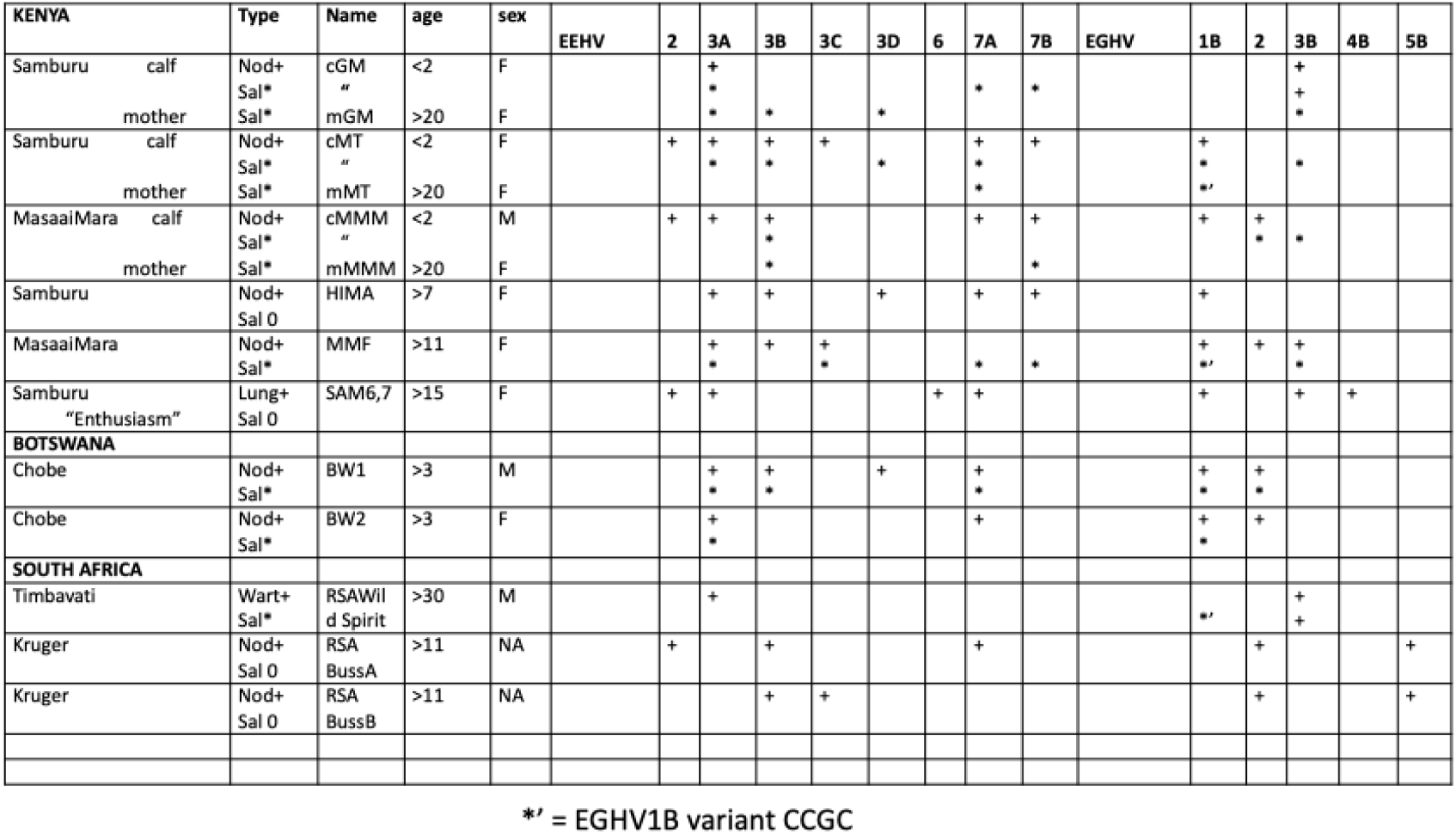
EEHV and EGHV in skin nodules (including matching saliva samples) and warts.

## RESULTS

### EEHV

DNA sequences of multiple species of EEHV and EGHV were identified in skin nodule biopsies from *L. africana* including one young female calf (Samburu NP, Kenya) and one young male calf (Maasai Mara NP, Kenya) with two nodules each; two young female calves (Samburu NP, Kenya and Chobe NP, Botswana, with one nodule each; and one young male calf with 4 trunk nodules (Chobe NP, Botswana); and 4 sub-adult females (2 Maasai Mara NP, Kenya and 2 Kruger NP, South Africa) with one nodule each; in ear wart-like lesions from an adult South African bull *L. africana*, (Timbavati, South Africa); and in a random field necropsy lung tissue (**Fig 1**) from a sub-adult female *L. africana* in Samburu, Kenya, (previously reported. [44]. DNA extracted from these samples were selectively screened using combinations of up to 21 different cPCR primers (including first, second and third round amplification for each when appropriate) that detected known selected high GC-rich EEHV3 or EEHV7 loci, as well as several more generic cPCR primer sets that also detected EEHV2 or EEHV6. 100% of nine (n=9) skin nodule biopsies (**Fig 3,4,5,6,7,8**) and one (n=1) ear wart biopsy (**Fig 9**) collected from wild *<u>L. africana</u>* adults and juveniles from Kenya (5), Botswana (2) and South Africa (2) were found to be low level positive by nested cPCR and Sanger sequencing for EEHV2, EEHV3A, EEHV3B, EEHV6, EEHV7A and EEHV7B and a wide variety of subtypes or strains, including novel subtypes of EEHV3B and EEHV7B and rare subtypes EEHV3C to EEHV3H as well as multiple examples of EGHV1B, EGHV2, EGHV3B, EGHV4,EGHV5B, EGHV3B, EGHV4B and EGHV5B. In a corollary study previously reported by Pearson et al, 2021, African Elephant Polyomavirus (AelPyV-1) was detected in these nodules. These findings suggest that further investigation is needed to ascertain whether infiltrating lymphocytes latently infected with EEHV and/or EGHV or other viruses such as AelPyV-1 may be responsible for such skin nodules on wild *L. africana*.[60]. **Fig 12**.

Saliva from Kenya and Botswana mother plus calf pairs (n=6, positive 6/6=100%) and from 26 (n=34, positive 29/40=73%) additional *L. africana* in Botswana, Kenya and South Africa and Zimbabwe were screened for GC-rich branch cPCR loci (POL, HEL, U71/gM, OBP, TK, vGCNT1 and UDG) yielding unambiguous EEHV DNA sequences of EEHV2, EEHV3A, EEHV3B, EEHV3C, EEHV 3D, EEHV6, EEHV7A and EEHV7B and additional unresolvable mixtures. Saliva samples from four of the same Kenya juveniles with nodules also detected between one and four subtypes of EEHV3A, EEHV3B, EEHV3C, EEHV3D and one or two subtypes of EEHV7A and EEHV7B as well as EGHV1B and/or EGHV3B each, some of which matched the strains in the skin nodules but mostly did not. Saliva from some mother/calf pairs were identical, but more often different.

It is important to note that many of these experimental primer sets (see Supplemental Information SI) were designed especially for this study, and often required multiple primer combinations including rounds 2A and 2B and rounds 3A and 3B to target specific EEHV genes. Additionally, many bright gel electrophoresis bands not of the expected base pairs for each cPCR round were sequenced in both forward and reverse directions in our attempts to distinguish between EEHV3A and EEHV3B, and EEHV7A and EEHV7B. Of the many thousands of likely postive gel electrophoresis bands generated of approximately the expected size, we present representative examples of our initial gels that yielded positive matches to EEHV, **Fig 14**, **Fig 15**, **Fig 16**. In lanes where no positive signal is shown, it is likely that the EEHV gene sequence targeted by the particular primer set may not have been present in the particular sample or another primer set combination would be required.

**Figure 14.**
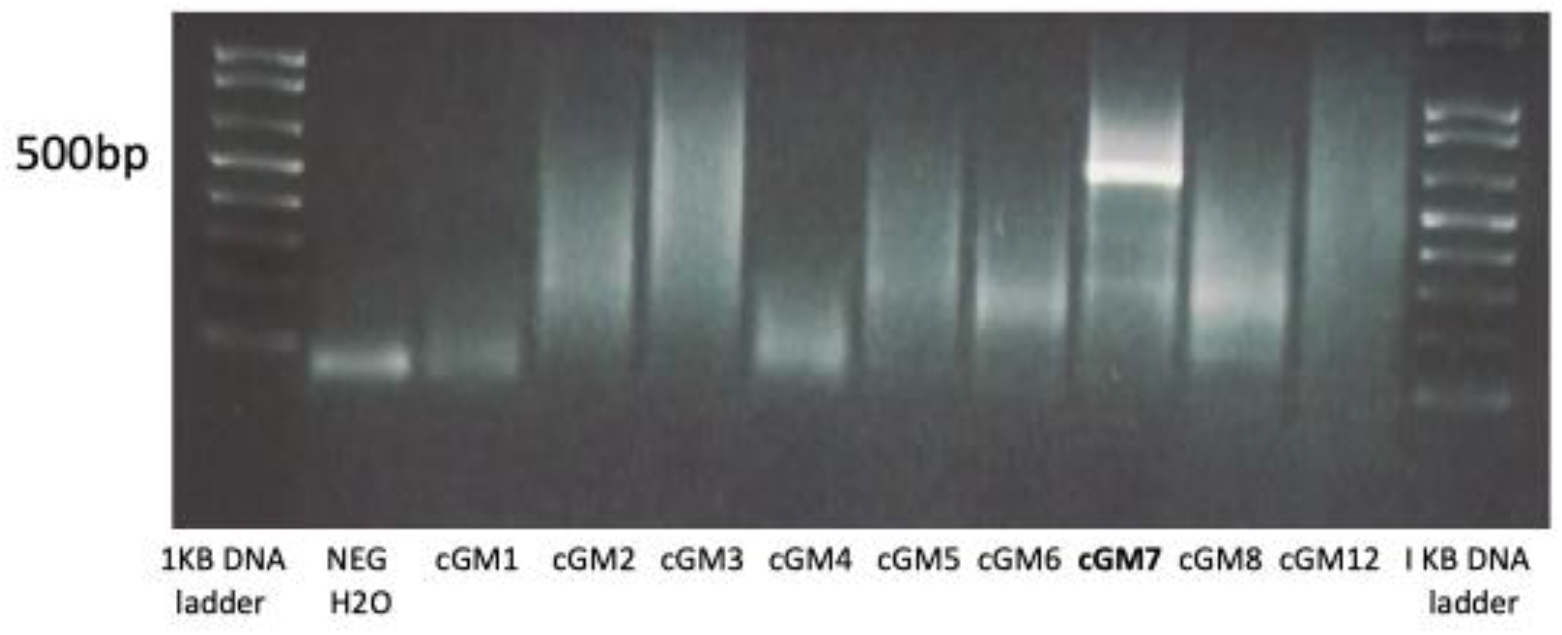
1.5% agarose gel of DNA extracted from KENYA calf cGM skin nodule biopsies (see **Fig 3**) and amplified by qPCR using Set 18 OBP rd2 primers 9394/9288, expect 345 bp. Bright band in Lane 9 (cGM7) of unexpected size was purified and Sanger sequenced yielding positive, unambiguous sequences for EEHV 3A. Imaged on BioRad ChemiDoc XRS Imaging System and photographed by Samsung Android cell phone. Genewiz Tracking #10-21977968.

**Figure 15.**
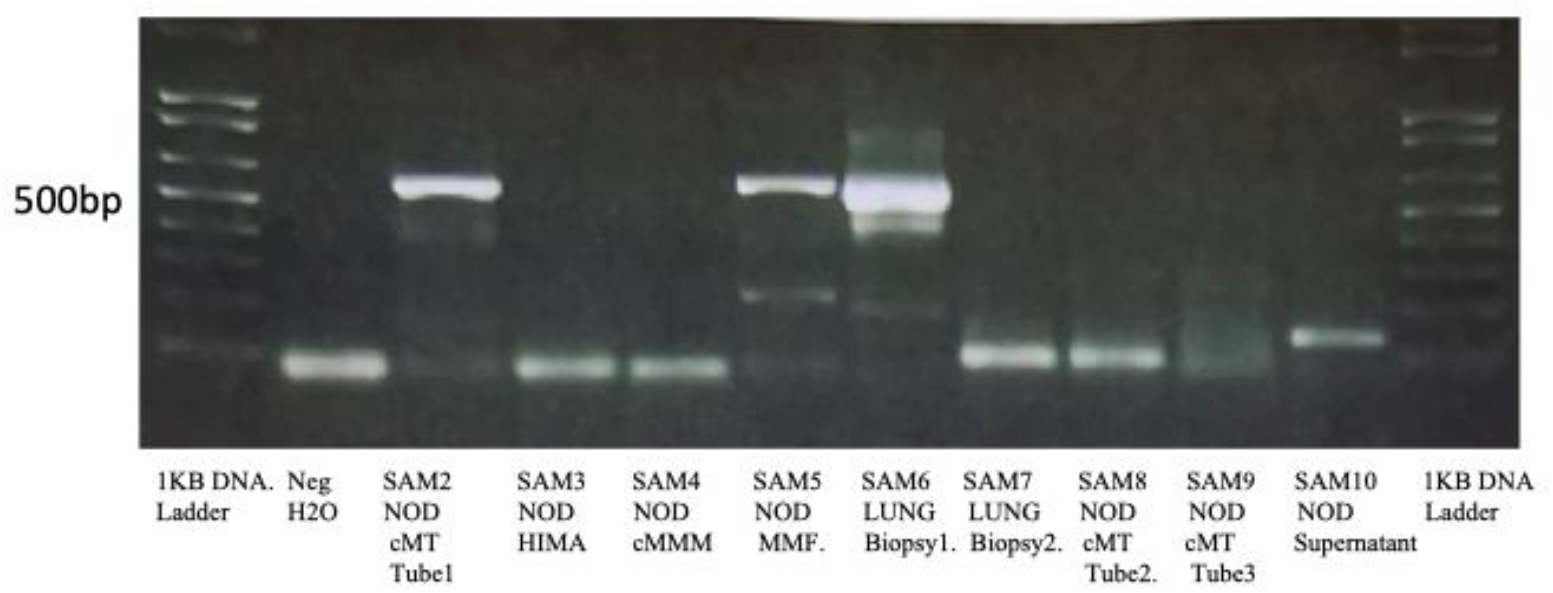
1.5% agarose gel of DNA extracted from skin nodule biopsies of KENYA calf cMT (see **Fig 3**)( SAM 2 NOD) HIMALAYA (SAM 3 NOD), calf MMM (SAM NOD 4), MMF NOD 5, (see **Fig 4**, **Fig 5)** and lung biopsies designated SAM 6 LUNG, SAM7 LUNG (see **Fig 1**) and amplified by qPCR using Set 18 OBP for EEHV3/4/7 rd2 primers 9394/9288, expected 345 bp. Bright bands of unexpected size were purified and Sanger sequenced to yield positive, unambiguous seqeunces for EEHV3B in skin nodules and EEHV3A and EEHV3B in lung biopsy. Imaged on BioRad ChemiDoc XRS Imaging System and photographed by Samsung Android cell phone. Genewiz Tracking #10-219228329.

**Figure 16.**
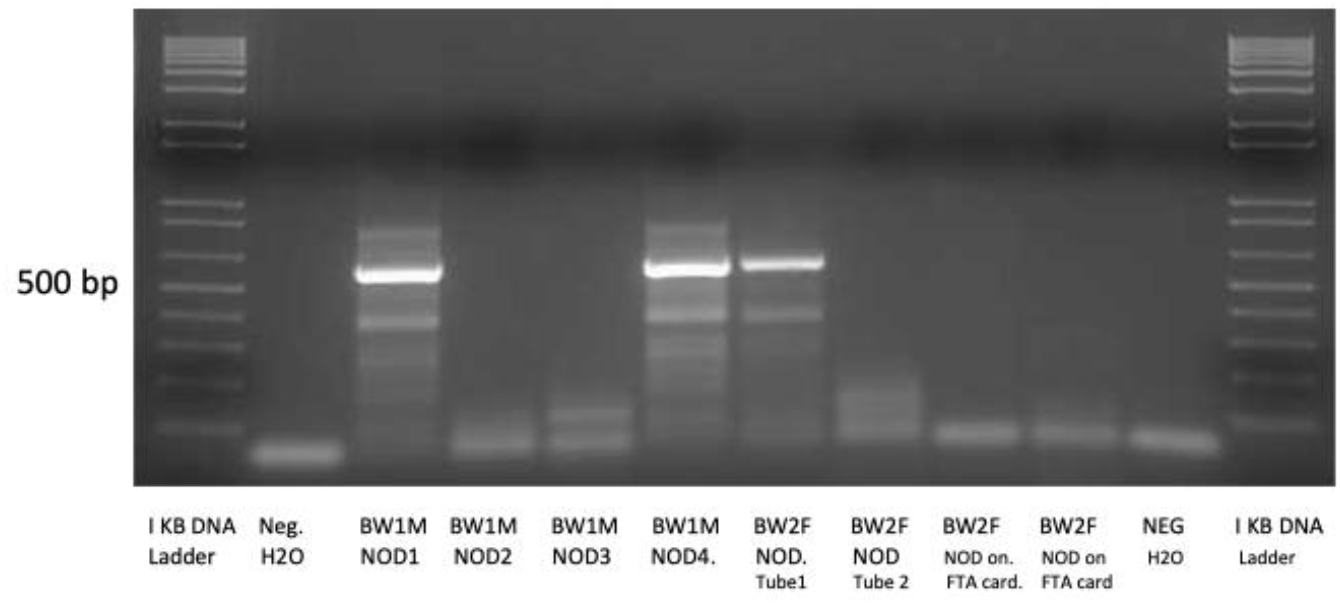
1.5% agarose gel of DNA extracted from skin nodule biopsies of BOTSWANA BW1M NOD 1 and BW1M NOD 4 and BW2F NOD (**see Fig 7**, **Fig 8**, **Fig 9**) and amplified by cPCR using Set 6B U77 HEL rd 2, primers R2-7=LGH7921/ L1=3198, expected 650 bp. Bright bands of expected size were purified and Sanger sequenced to yield positive, unambiguous seqeunces for EEHV 3A in BW1M and EEHV7A in BW2F. Imaged on BioRad ChemiDoc XRS Imaging System and photographed by SamsungAndroid cell phone. Genewiz Tracking #10-237155216.

Saliva swabs from 25 *L. cyclotis* were screened by cPCR and Sanger sequencing with more than six GC-rich branch EEHV primer sets including POL, HEL, TER, UDG, TK and OBP loci. Over half of 25 saliva samples from *L. cyclotis* in Gabon were positive for at least one EEHV*3* or EEHV7 strain, and once for a novel EEHV6, but no EEHV2 was detected, and many of these were also present as unusual, rare variants of non-A/non B subtypes of EEHV that we found in nodules and saliva from *L. africana.* In terms of number of distinct virus strains in individual samples, five were negative, 8x had just one strain, 8x had two strains, 3x had three strains, 1x had four strains and 1x had five different strains present. Unusually, unlike the results from the skin nodules, only a single EEHV strain was detected in any one Gabon saliva sample. The high prevalence of EEHV3B suggests that most likely EEHV3B and the more rare EEHV3C, EEHV3D, EEHV3E, EEHV3F subtypes have been exchanged and recombined during overlapping range interactions between *L. africana* and *L. cyclotis* hosts. These results closely resemble previous published findings by Zong et al 2015 of EEHV in lung nodules and necropsy lung tissue [44]. We attribute the lower percentage of positives in the Gabon samples (Gabon positive 13/25=52%) as compared to samples from Kenya and Botswana to insufficient volume of saliva collected. in the swabs, **Fig 17**.

**Fig 17.**
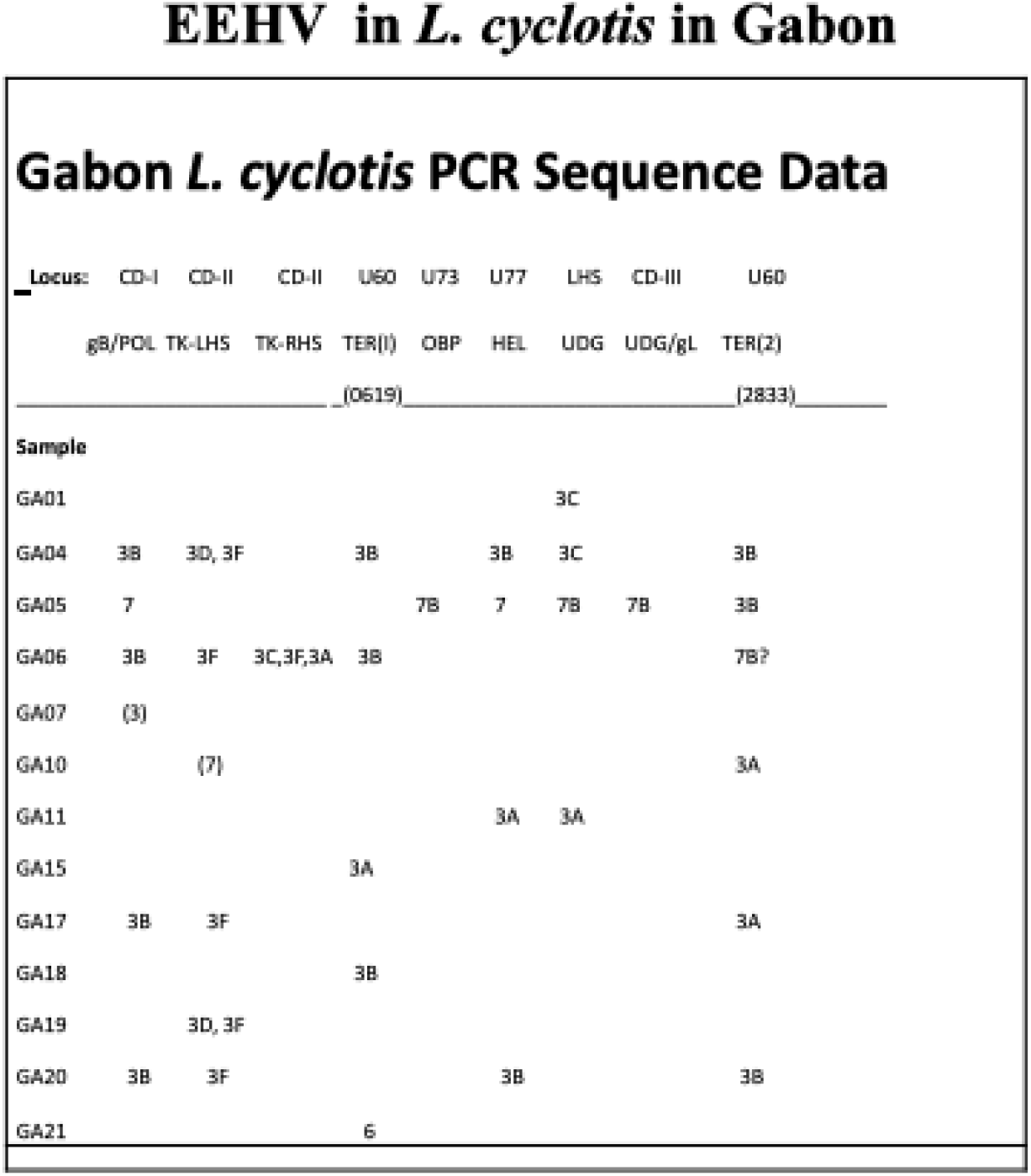
EEHV found in *L. cyclotis* in Gabon

**Fig 18.**
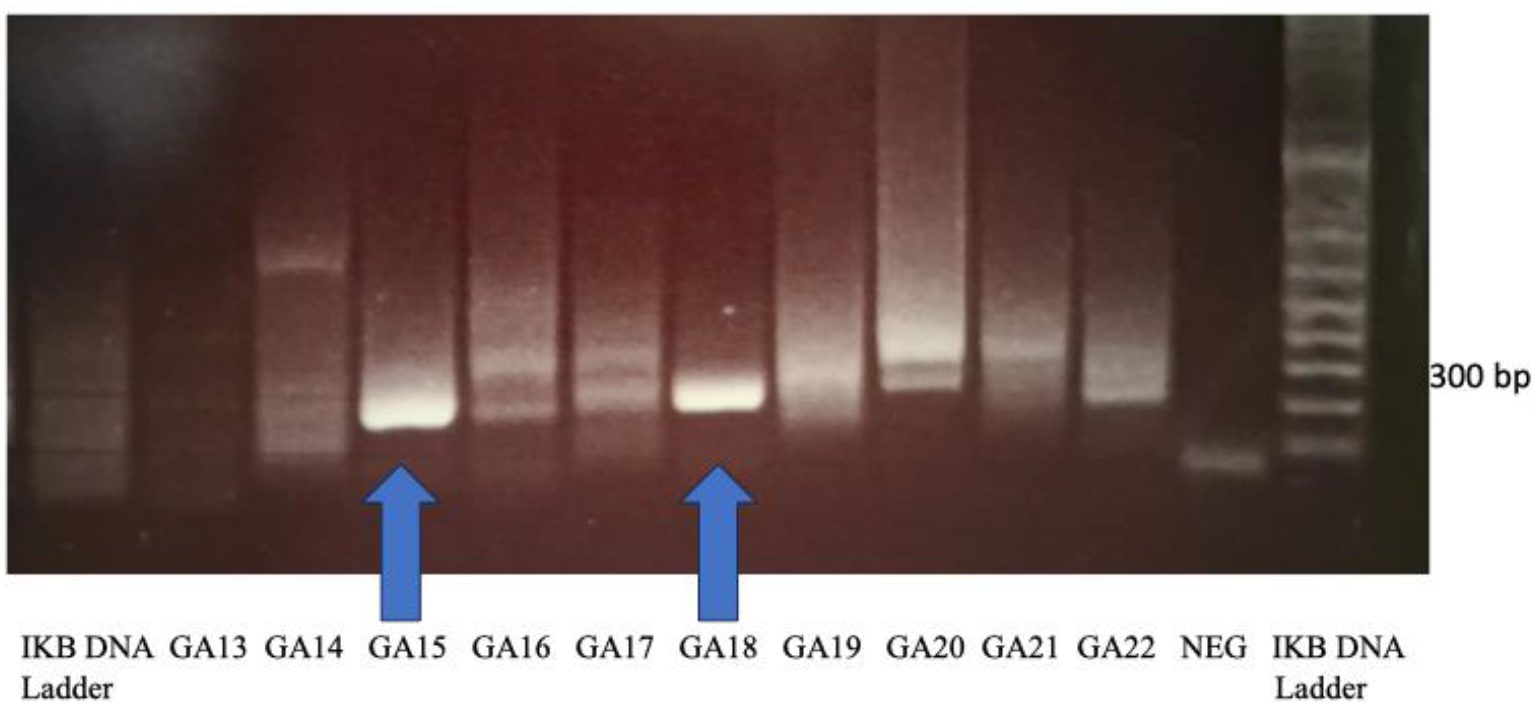
1.5% agarose gel of DNA extracted from GABON saliva samples and amplified by cPCR using Set7 HEL rd2B, Primers R2-7/L0-7, expected 250 bp. Arrows indicate bright bands of expected size that were purified and Sanger sequenced to yield positive, unambiguous seqeunces for EEHV 3A (GA 15)and EEHV 3B (GA 18). Imaged on Clare Chemical Research Dark Reader Tansilluminator and photographed by Samsung Android cell phone. Genewiz Tracing #30-194890619.

Our temporal saliva study over seven years of seven *L. africana* at Six Flags Safari Park, NJ, USA, showed that EEHV2, EEHV3A, EEHV3B, EEHV6, EEHV7A and multiple EGHVs are periodically shed in saliva, with frequently more than one virus being detectable at a time.

Primers used were TER, POL, OBP and UDG. Saliva from two of seven adult females, SFL and SFJ, were collected approximately bi-weekly over a full 12-month time period from January, 2012 through January, 2013 for a total of 43 samples each for Year One (Y1) and thereafter 12 times until through 2019 for a total of 55 samples each. The other five herd members were sampled twice during the 2012 and thereafter a total of twelve times for a total of 14 samples each, for an overall total of 180 total saliva samples. Examples of gels are shown, **Fig 19**, **Fig 20**.

**Figure 19.**
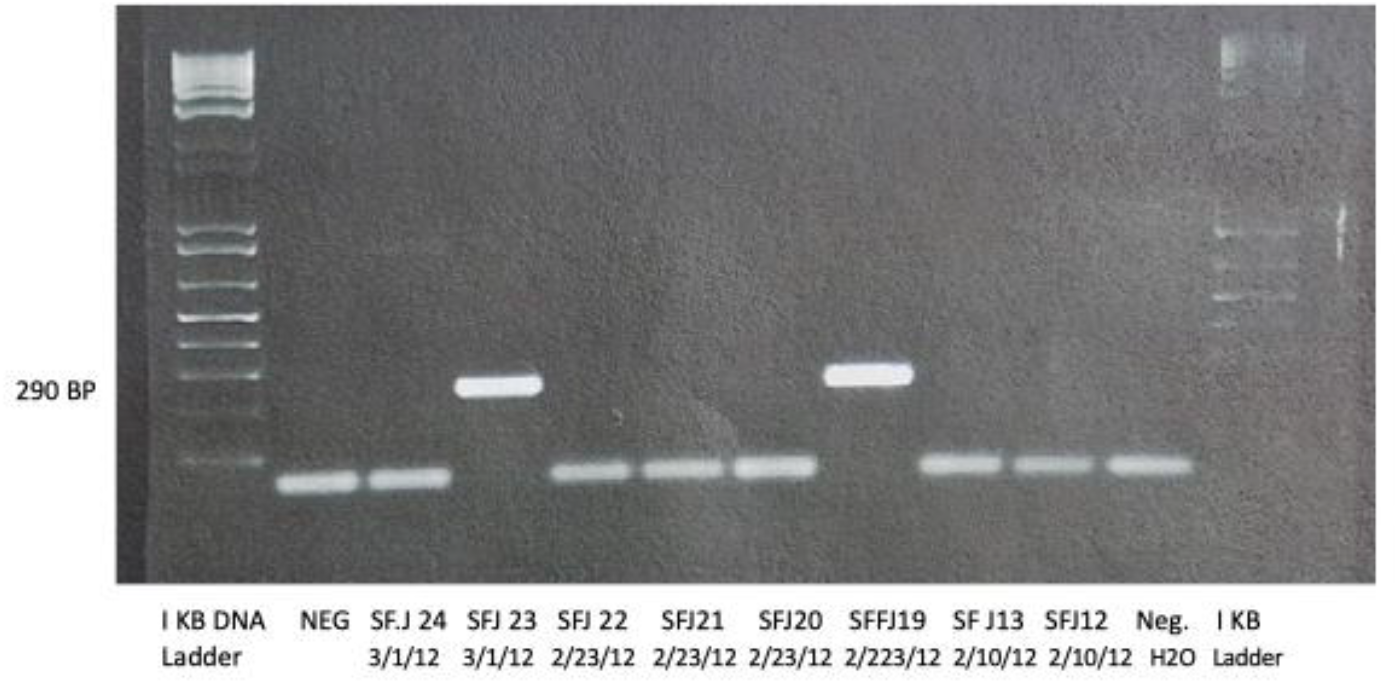
1.5% agarose gel of DNA extracted from SIX FLAGS SFJ saliva samples (see **Fig 10)** and amplified by cPCR using Set 2A TER rd2 primers 3025B/7576B, expected 290 base pairs. Multiple swab collections were taken initially on the same days as *L. africana* SFJ became used to saliva collection. Bright bands of expected size that were purified and Sanger sequenced to yield positive, unambiguous seqeunces for EEHV 2 in SFJ23 and SFJ19 Imaged on BioRad ChemiDoc XRS Imaging System and photographed by Samsung Android cell phone. Genewiz Tracking #10-211611514.

**Figure 20.**
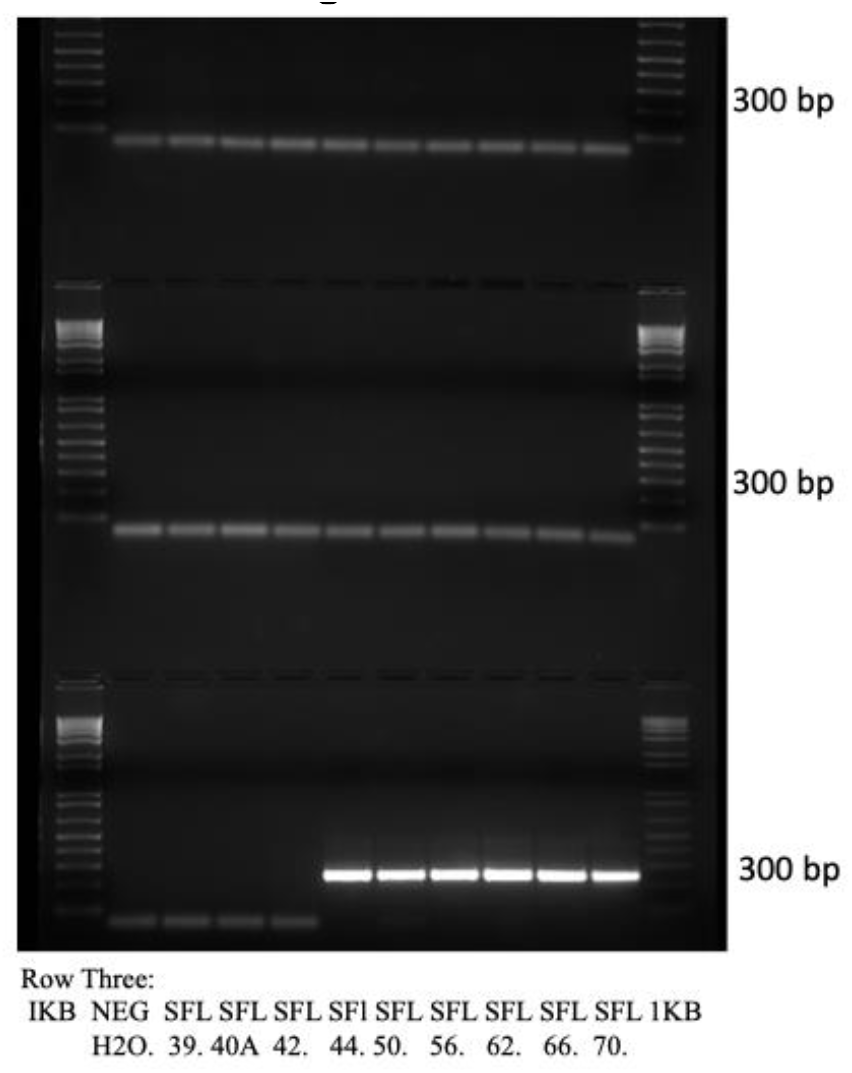
1.5% agarose gel of DNA extracted from SFL saliva samples, amplifiedd by cPCR using Set2A TER rd 2, primers 3025B/L2-2A expect 290 bp. TOP ROW: Lanes 3-10 (SFL#1 #2,#3,#4,#9,#10,#11,#15,#16, collected 1/27 to 2/23/2012) = negative); MID ROW: lanes 3-10 (SFL#17 #18,25,27,29,31,33,35,37, collected 2/23 to 5/2/2012 = negative; BOTTOM ROW: lanes 3-10 (#40A,42,44,50,56,62,66,70, collected 5/8 to 7/15/2012) only lanes 6-11=EEHV6. Bright bands of expected size were purified and Sanger sequenced to yield positive, unambiguous seqeunces for EEHV 6 in SFL #44 ((5/24/2012) through SFL#70 (7/15/2012), indicating no shedding of EEHV 6 before 5/24/2012; however shedding of EEHV6 continued sporadically through 9/27/2019 (not shown). Imaged on BioRad ChemiDoc XRS Imaging System and photographed by Samsung Android cell phone. Genewiz Tracking #212152244.

Overall, SFJ scored positive for EEHV2 in 17/43 = 40% of Y1 samples while SFL scored positive for EEHV2 in 8/43=19% of Y1 samples. In contrast, SFJ was positive for EEHV6 in 17/43=40% Y1 samples and SFL was positive for EEHV6 in 15/43=35% of Y1 samples. Both SFL and SFJ shed both EEHV3A and EEHV3B intermittently for a total of SFJ 11/ 43=26% and SFL 10/43=23%. SFJ was positive four times for four different viruses at the same time (EEHV2, EEHV3A, EEHV3B and EEHV6) whereas SFL was positive at some point for two or three different EEHV species at the same time (either EEHV2, EEHV3B and EEHV6 or EEHV2, EEHV3B). Overall, among seven adult *L. africana* tested at Six Flags Safari Park from 2012 to 2019, 6/7=86% were positive for EEHV2, 5/7 = 71% were positive for EEHV6, 4/7=57% for EEHV3B, 4/7=57% for EEHV3A and 2/7 = 29% were positive for EEHV7A. Only a single sample from SFJ=EEHV2 POL (SFJ sample #4, January, 2012), gave a band of sufficient intensity to obtain clear unambiguous DNA sequence data directly from the first round cPCR amplification, **Fig 21**.

**Fig 21.**
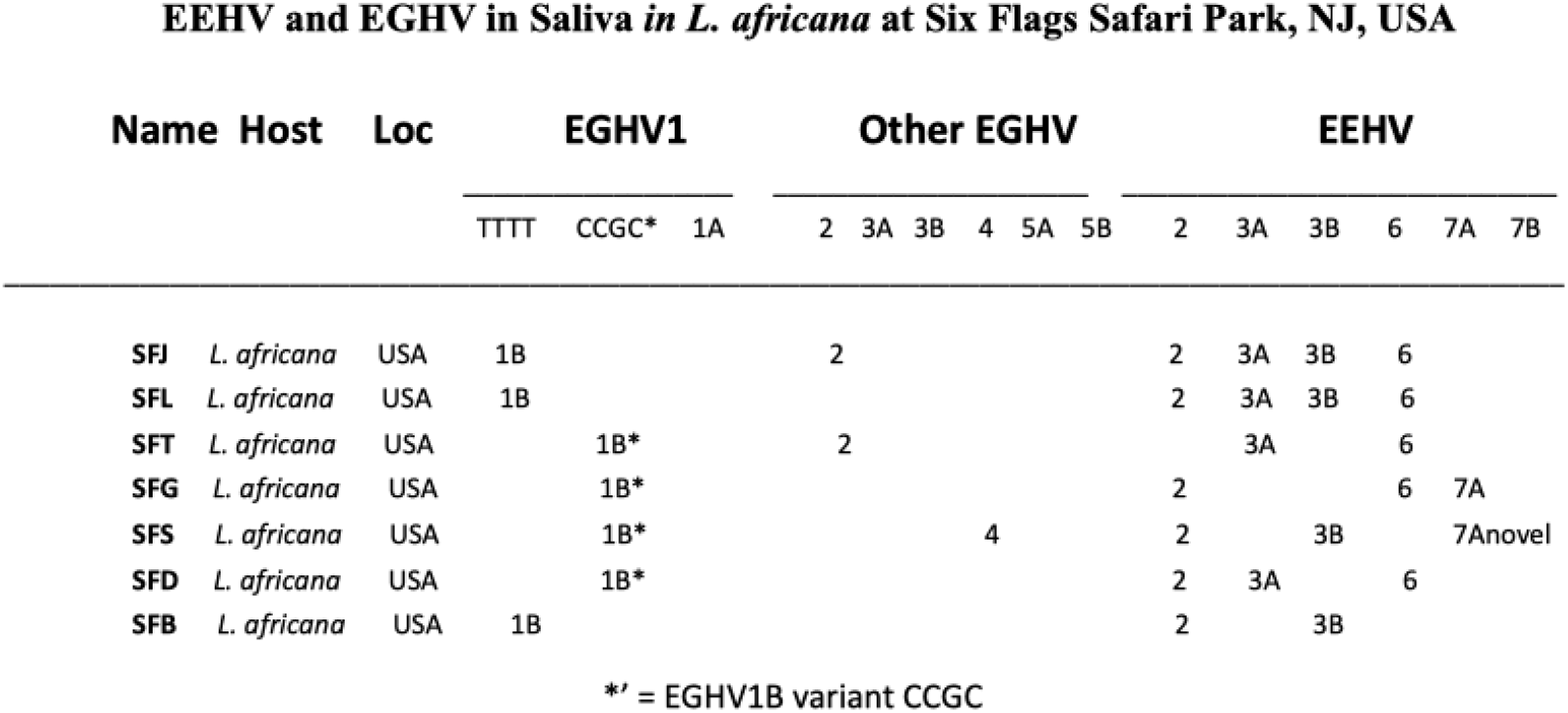
EEHV and EGHV in saliva from *L. africana* at Six Flags Safari Park, NJ, USA. SFJ saliva sample #97 collected on November 21,2012 has been designated the prototype for EGHV1B, (NAG 9, GenBank #KC810011.1).

### EGHV

In addition to the multiple EEHV subtypes associated with EHD, most healthy asymptomatic adult elephants harbor and periodically shed DNA from the second herpesvirus sub-family EGHV. Overall nodule biopsies from six of the seven wild *L. africana,* the lung biopsies, South non-nodular ear wart biopsy and more than half of the saliva samples from 29 wild *L. Africana,* 25 *L. cyclotis* and 70 *L. africana* from North America proved positive for clean unambiguous detection of at least one type of EGHV, although many more clearly contained unresolvable mixture of multiple EGHV genome types.

Sixteen examples of **EGHV1B** (the most abundant EGHV species we encountered) were detected within sub-Saharan Africa, including two invariant polymorphism patterns that differ by only 4/450-bp (0.9%). The more commonly found TTTT pattern was seen in 13 of 14 EGHV1B cases from *L.africana* within Africa, while the more rare CCGC pattern was found only once in *L. africana* adult bull Matambu (SAG5, Timbavati, South Africa) in continental Africa. All three EGHV1B positive samples from *L. cyclotis* fall into the same CCGC variant suggesting distinct host species origins for these two variants. These same two polymorphic variants were detected in three whole herd saliva sets from Six Flags Safari Park where SFJ, SFL, herd mates SFG, SFS and SFB carried the EGHV1B TTTT variant whereas herd mates SFT and SFD carried the CCGC variant that was identical to South African RSA Matambu (see **Fig 21)**. In saliva samples from additional USA *L. africana* (not shown) we found only the CCGC variant in less than one-quarter of the samples tested. **Fig 22**, **Fig 23**.

**FIG 22.**
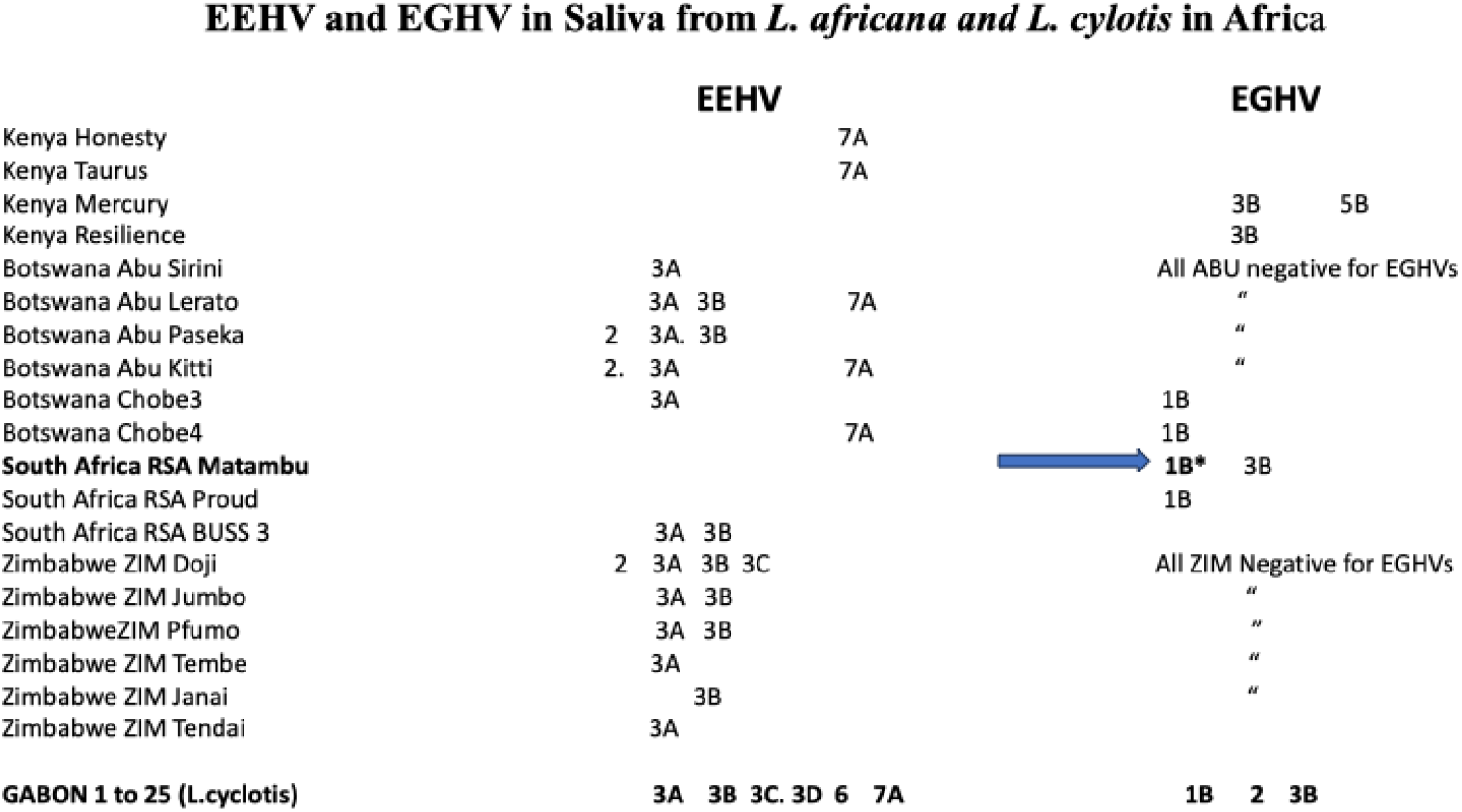
EEHV and EGHV in saliva from *L. africana and L. cylotis* in Africa. *= EGHV1B variant CCGC, all others = EGHV 1B variant TTTT.

**Fig 23.**
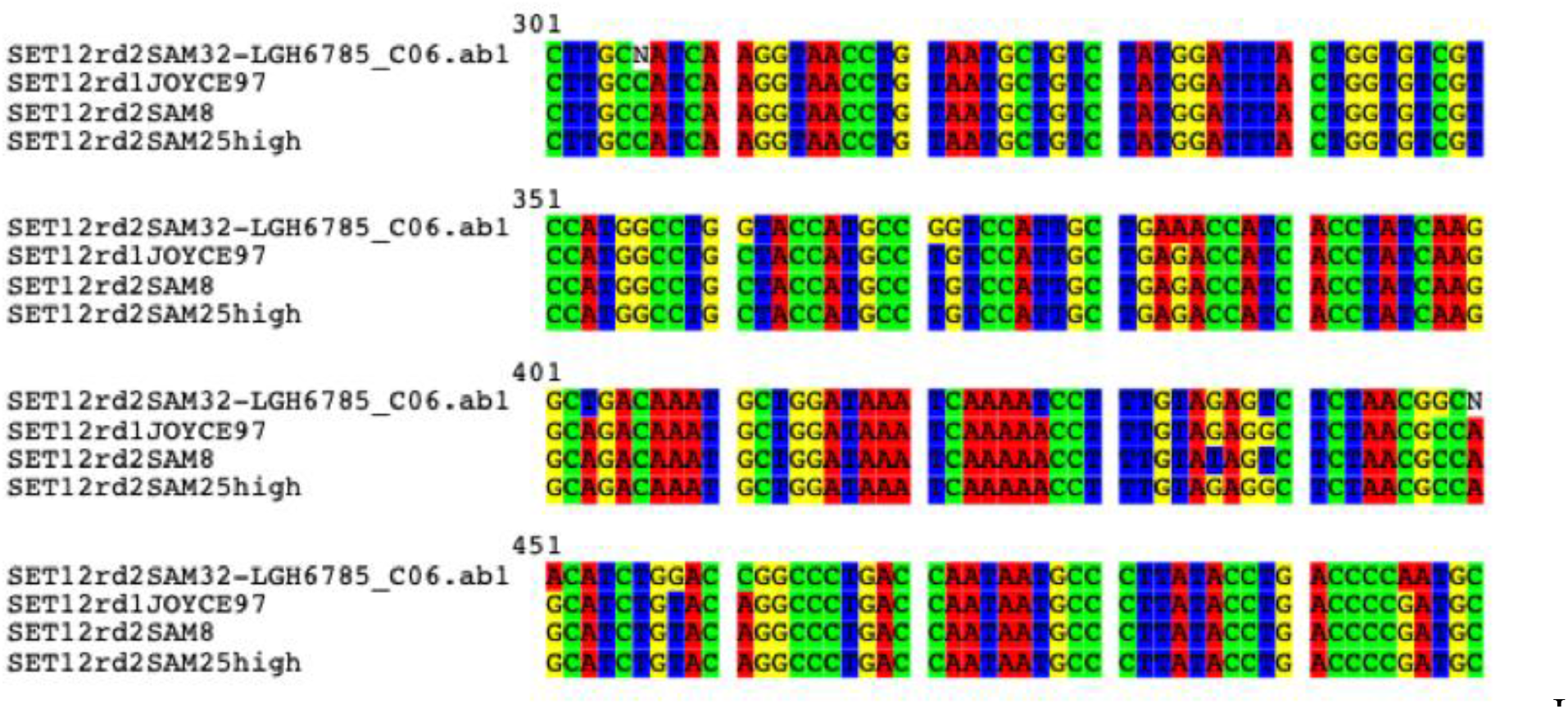
DNA alignment for EGHV1B exhibiting distinct polymorphisms among saliva and skin nodules: SAM 32= cMT saliva; SFJ #97= saliva (prototype For EGHV1B); SAM 8 cMT NOD, and SAM 25=cMMM NOD

**EGHV2** was detected in POL locus in skin nodules from wild *L. africana* RSA BussA, KENYA MMM, KENYA MMF, and Botswana BWM1, BWF2 and in saliva from Six Flags Safari Park SFJ and SFT which are identical to those EGHV2 sequences reported in a 38 year-old zoo *L. africana,* GenBank acc# KU726836, # KU726837 [35] and were just one to four bp different from the prototype *E. maximus* EGHV2 POL.

**EGHV3B** was detected frequently in skin nodules, ear warts and saliva from *L. africana* and in saliva from *L. cyclotis*. Quite unexpectedly however, we detected widespread presence of EGHV3B in necropsy organs from an adult female *L. africana,* NAG31, in a USA zoo, including in ear lesions, heart, lymph node salivary gland, serum and thymus. We have rarely detected any EGHVs in any internal organ tissues from *EHD* cases or necropsy samples in USA zoos. Surprisingly, no African Elephant Polyomavirus DNA was detected in the ear lesions.

**Figure 24.**
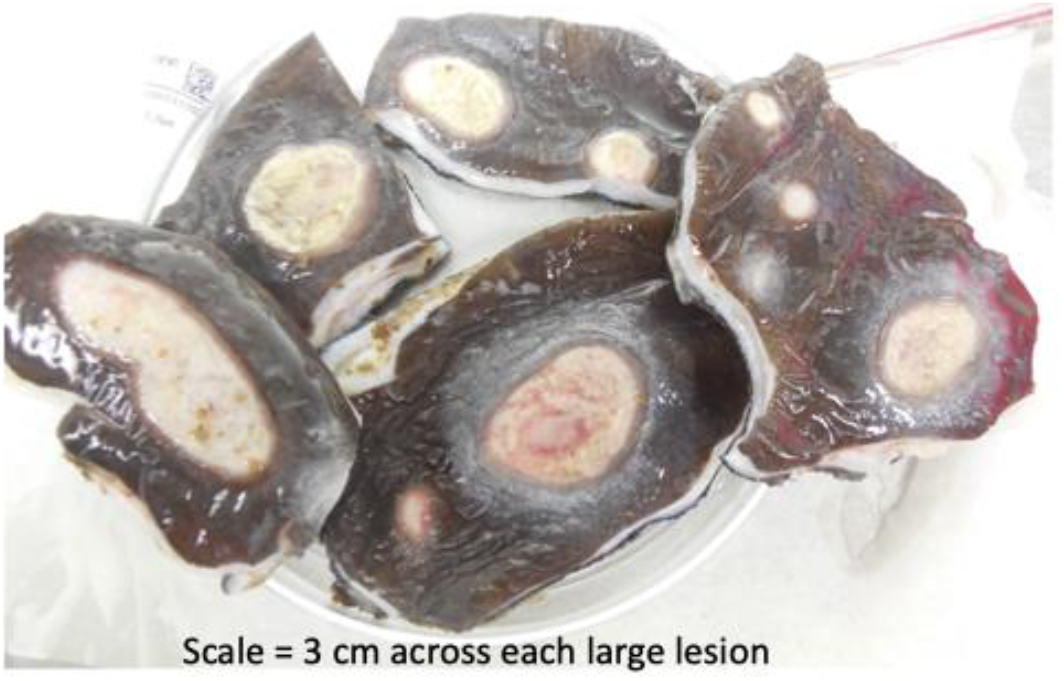
Ear lesions on back of ear from necropsy samples of adult female *L. africana* NAG31 in USA zoo, received by Pearson, 2014. Ear lesions, heart, lymph node salivary gland, serum and thymus were positive for EGHV3B only, no EEHV was detected, credit Pearson, 2014.

**EGHV4** was detected in saliva from SFS at Six Flags Safari Park; in the young zoo *L. africana* that survived EEHV3B [35]; and in wild *L. africana* lung biopsies designated KENYA Sam 6 and Sam 7(see **Fig 1**) collected by Pearson in 2011. [44, 47–50]. Subsequently, we discovered an evidently novel Asian elephant version of EGHV4 in spleen tissue from a three year old male captive-born *E. maximus* NAG72 in New Mexico, USA, that died suddenly of EHD, GenBank Acc. #PQ379918. Identical 250 bp sequences were obtained from saliva from a captive-born living male juvenile *E. maximus* NAG71 in Texas and in the spleen from a deceased elderly USA *E. maximus* NAG56 in Florida, USA. These results led us to reassign all the previous *Loxodonta-*associated EGHV4s and *Elephas*-associated EGHV4*s* as distinct subspecies EGHV4A or EGHV4B endogenous to either *Elephas* or *Loxodonta* respectively.

**EGHV5B** was detected in skin nodule biopsies from two sub-adult *Loxodonta africana* in Kruger National Park, South Africa: BussA SAG3, BussB SAG4, and in saliva from an adult femal*e L. africana* in Kenya designated KENYA Mercury KG16. In this case, the 450-bp DNA sequences obtained after second round PCR with the generic Pan-EGHV primers were identical to one another and to those from the prototype EGHV5B (GenBank accession #JX268522. [55]. Most recently, Sanger sequencing from first round cPCR using EGHV PAN POL primers R1/L1 of a skin nodule biopsy from the trunk of a 3 year-old captive born *L. africana* in England yielded a 439 base pair sequence of EGHV5B, GenBank Acc# identical to the prototype NAG8. The sequence obtained from the R1 and L1 primers, was round 1 PCR, R1 LGH6784B 5’-GTGGTKGACTTTGCYAGCCTSTACCC-3’ L1 LGH7489 5’-GTCRGTGTCYCCGTAGAYNAC-3’. This is the first time we have been able to compare EGHV sequence results from a skin nodule on a *L. africana* residing outside of Africa to skin nodule biopsies and saliva collected in Botswana and Kenya.

**Figure 25.**
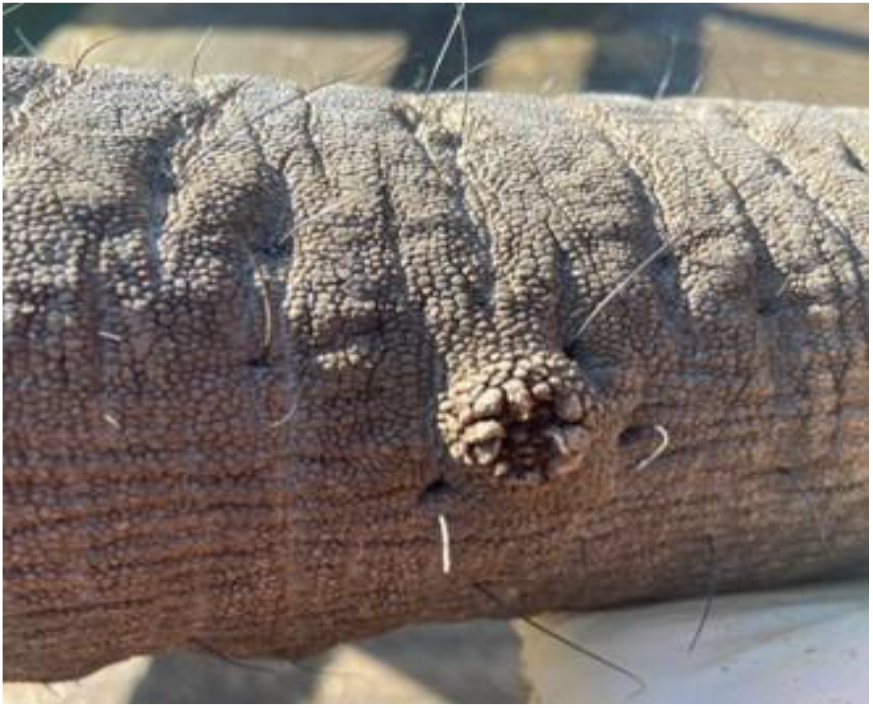
Trunk nodule on 3 year old *L. africana* positive for EGHV5B, credit J. Hopper, DVM, Howletts Wild Animal Trust; DNA extraction, cPCR and Sanger sequencing provided by A. Dasjerti, DVM, APHA, UK, for Howletts Wild Animal Trust, England, 2024.

## DISCUSSION

EEHV is an ancient cladal group that evolved along with the ancestors of modern proboscidean hosts for tens of millions of years. They form two major branches with either AT-rich versus GC-rich character, which in turn split into seven distinctive species that diverge overall from each other by at least 16 to 20% at the nucleotide level. Each species also displays A plus B subtypes that may contain up to ten localized non-adjacent chimeric domains (CD) or hypervariable gene blocks encompassing up to seven distinct subtype clusters (**Fig 26**) with large (15 to 50%) protein level differences [61–65]. The five EGHV species POL genes are widely diverged displaying between 37 and 45% nucleotide polymorphisms and fall into three ancient deep lineages within the mammalian gamma herpesviruses corresponding to estimated last common ancestors (LCA) occurring about 200 million years ago [66–69]. Four of the EGHV species (EGHV1, EGHV3, EGHV4 and EGHV5, but not EGHV2) further divide into consistent A and B diasporas that differ by around 3 to 5% (15 to 24/450-bp polymorphisms) and are largely specific for *Elephas* versus *Loxodonta* hosts respectively [70,71]. **Fig 26**.

**Fig 26.**
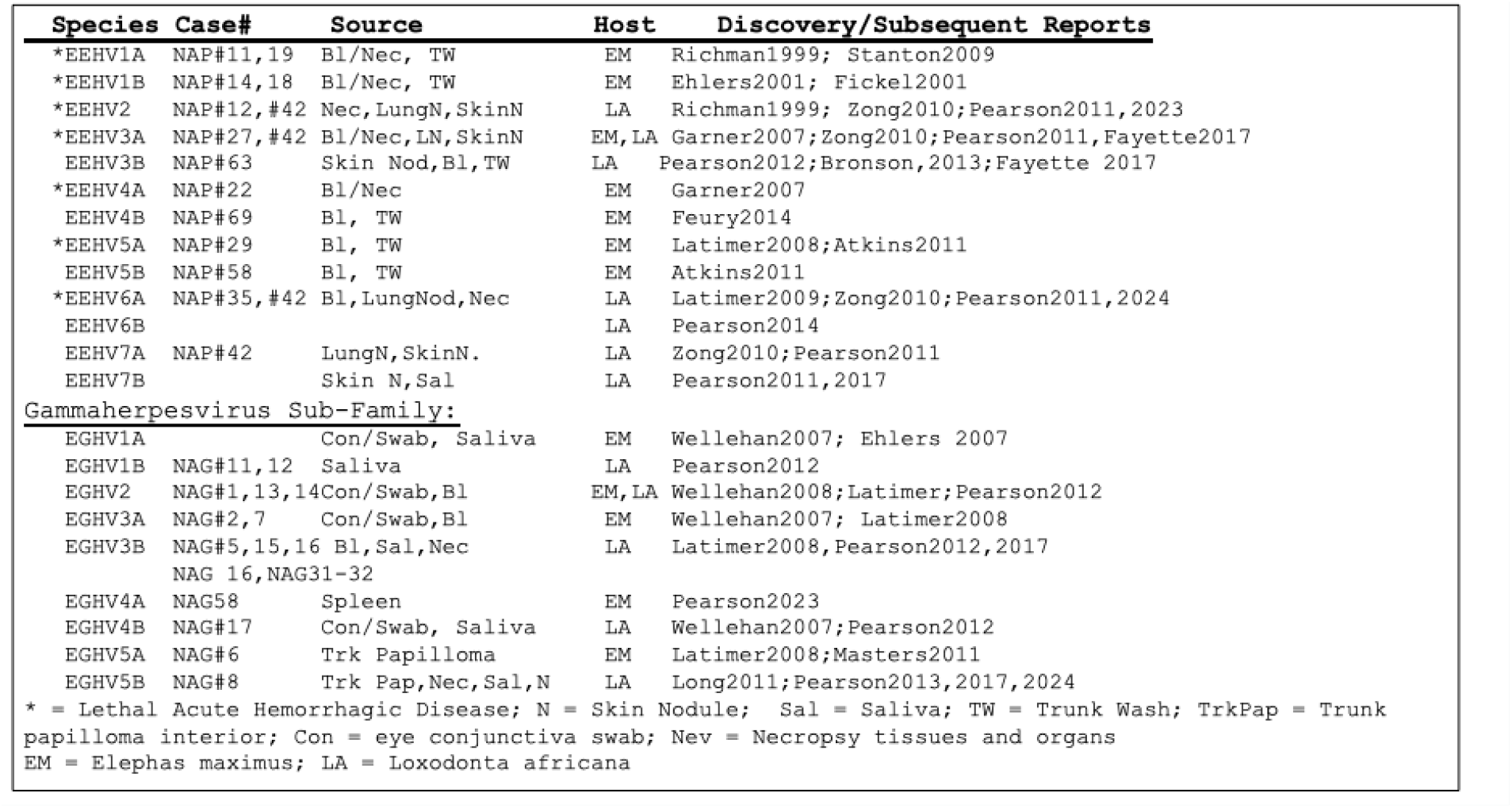
EEHV and EGHV Species and Subtypes.

Nearly all skin nodules examined proved to be much more complicated, contained multiple different virus types and strains and gave at least one unambiguous positive result with one or more of the primer pairs used. Furthermore, a large fraction evidently harbored multiple EEHV genome types and indeed unlike with any Asian elephant EHD viremic trunk wash or saliva swab samples evaluated in other studies, many of the skin nodules gave mixed DNA sequences (see **Fig 12**) from multiple different genomes being present within the same purified PCR bands.

This phenomenon of multiple simultaneously infecting genomes that sometimes gave mixed DNA sequence profiles was also seen in the previously reported study by Zong et al, 2015 [44] with the much smaller number of African elephant lung nodules, but it has been exclusive to just the skin and lung nodules or to asymptomatic necropsy samples, and has been very rare or non-existent within the saliva studies even from the same group of elephants. Note that some mixtures gave clean enough sequences that they could be resolved with or without additional nested PCR analysis, although many could not, and the latter have just been recorded here as “mixed 3/7” or “mixed 3A/3B” or “multiple 3A” strains present, without accompanying GenBank DNA sequence files. However, wherever possible clean DNA sequence files if of sufficient length to reveal phylogenetically useful characteristic polymorphisms have been generated, edited and submitted to GenBank.

A curious feature of the results from the Kenya and Botswana skin nodules is that no two nodules from the same animal seemed to have the same complement of virus species, subspecies or strains present and that the strains even of say just EEHV3A present in different nodules from the same animal often proved to be distinctly different with multiple novel polymorphisms present. Indeed, for EEHV3 especially, the large number of nucleotide variations found between the numerous different strains encountered and patterns of subgrouping among them have often allowed us to designate more than just EEHV3A and EEHV3B and further define subtypes EEHV3C, EEHV3D and EEHV3E that may contain up to ten localized nonadjacent chimeric domains (CD) or hypervariable gene blocks encompassing anywhere up to seven distinct subtype clusters with large (15 to 50%) protein level differences. EEHV3 strains fall into two major chimeric subtype clusters similar to those described previously in *Elephas* as EEHV1AB, EEHV4A/B and EEHV5A/B. [53]. In EEHV3A strains only, parts of the CDs including vECTL, gB. gH, TK, vGPCR1, gL and UDG can instead have several other alternative highly diverged but unlinked subtype clusters (designated EEHV3C, EEHV3D, EEHV3E, and EEHV3F, (GenBank acc.# MN373268, NC077039, NAP42 EEHV7A GenBank acc# KU321582, NAP62 Samson EEHV3B KT832467.1-KT832476.1, KT832491.1-KT832495.1. These differ greatly (by between 15 and 45%) in several linked but nonadjacent chimeric domains (CD-I, CD-II, CD-III, and CD-V) totaling 15 to 20-kb in size, but by just 2 - 3% everywhere else. Our ongoing genetic evaluation of the recent EHD cases in *L. africana* attributable to EEHV2 and EEHV6 suggest that similar division into A and B subtypes may be appropriate, as found in EEHV3A/3B and EEHV7A/7B pairs that differ by 7 - 9% overall with many other genes not just the CDs varying significantly. (GenBank acc# NC077039, MN373268). [51,52]. Based largely on the summed higher levels of individual protein divergence across the whole EEHV3A versus EEHV3B genome as well as the relatively high frequency of EEHV3B being found in saliva swabs from Gabon, compared to EEHV3A being far more prevalent in Botswana, Kenya, South Africa and Zimbabwe, we judge that these two EEHV3 groups should in fact be designated as distinct species with EEHV3A likely having evolved in *L. africana* and with EEHV3B probably having evolved originally in *L. cyclotis*.

The figures presented here illustrate several examples of the high levels of divergence that have been found to be exhibited at selected PCR gene loci by multiple species, subtypes and strains across the entire spectrum of the GC-rich branch of *Probosciviruses*. **Fig 27** shows a phylogenetic tree for the U48.5(TK) protein comparing 17 examples of EEHV3 and three of EEHV7 identified in biopsied skin nodules or saliva swabs collected from healthy juvenile and adult from *L. africana* in Kenya, South Africa, Botswana and North America, as well as three from *<u>L. cyclotis</u>*in Gabon (*). The prototypes for EEHV4A(Jennie) and EEHV4B(Baylor) from *E. maximus* are also shown. Note that four different samples for EEHV3B (Samson) [35] from whole blood, trunk wash, and saliva and determined in different laboratories were identical, but those classified as EEHV3 formed five distinct clusters designated 3A, 3B, 3C, 3D and 3E subtypes. Similarly, **Fig 28** gives a DNA level phylogenetic tree for the U71(gM) locus comparing 26 positive examples of EEHV3 or EEHV7 skin nodule or saliva swab DNA from *L. africana*, as well as the two prototype EEHV4 versions. At this locus both the EEHV3s and the EEHV4s split into just the classic 3A versus 3B and 4A versus 4B diasporic patterns. The genomes from all four major EEHV species found in African elephant hosts have now been completed with annotated versions of the prototype EEHV2A, EEHV3A and EEHV6 strains all available at GenBank, and with the EEHV3B and EEHV5B data submitted but not yet released. [72]]. All raw EEHV and EGHV sequence data generated by Pearson while at Princeton University and Fox Chase Cancer Center are accessible in digital format through The International Elephant Foundation.

**Figure 27:**
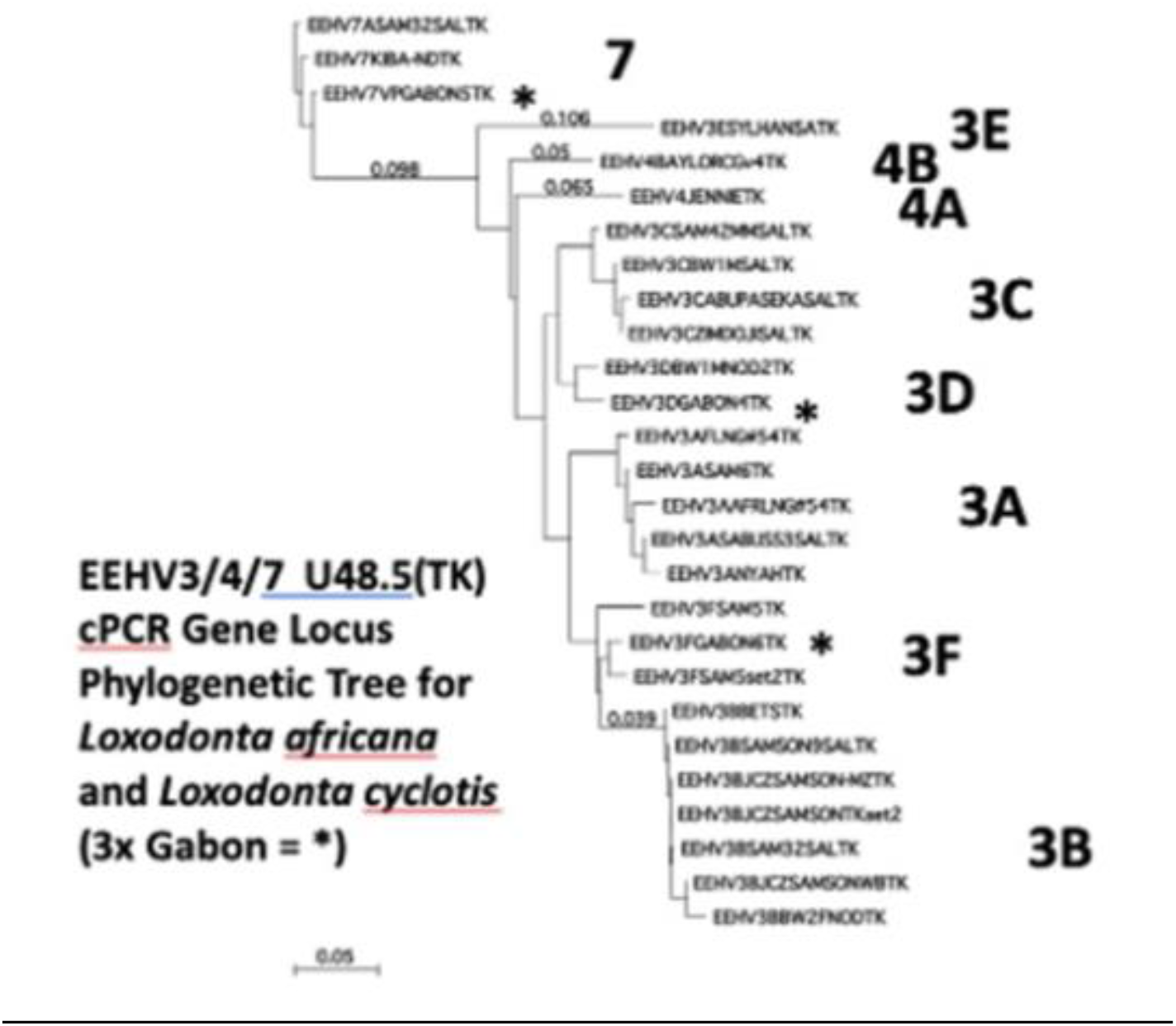
DNA level phylogenetic tree showing species and strain subtyping divergence at the U48.5(TK) PCR locus among 17 strains of EEHV3 and three of EEHV7 from *L. africana* and *L. cyclotis* compared to the prototypes of EEHV4A and EEHV4B.

**Figure 28:**
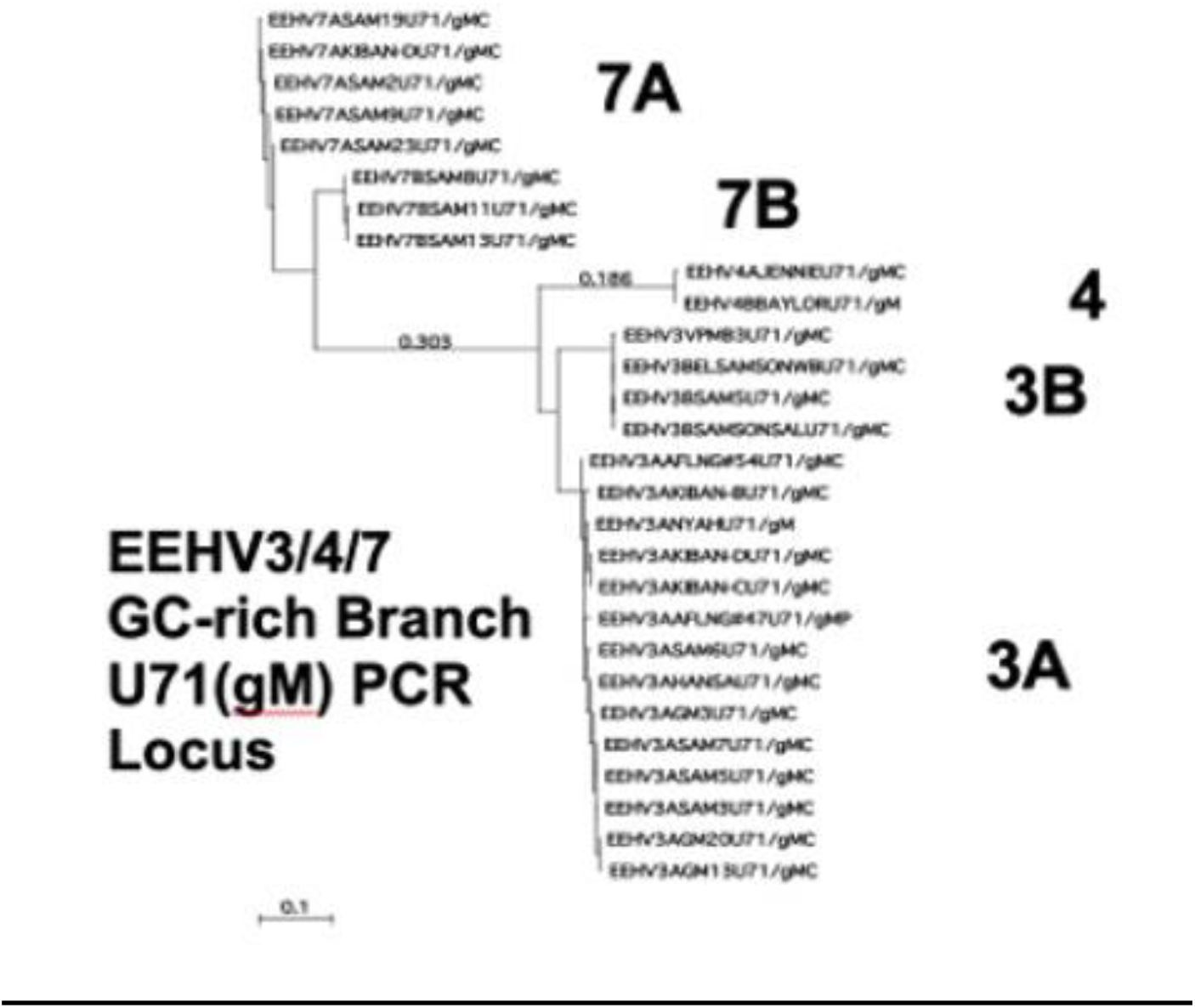
DNA level phylogenetic tree for the U71/gM PCR locus for 26 examples of the EEHV3 or EEHV7 species found in the GC-rich branch of the Probosciviruses from *Loxodonta africana* compared to the prototype *E. maximus* versions of EEHV4A(Jennie) and EEHV4B(Baylor).

It is important to note that in our saliva swab analyses, as opposed to skin nodules, very few samples were positive directly on first-round cPCR, and most were only detected after either second-round or even third-round nested cPCR. these represent values of only a few 100 to a few thousand viral genome equivalents (vge) per ul/ug of DNA. In contrast, in the blood or tissue from severe EHD cases the cPCR is usually still first-round positive even after 10,000-fold dilution of the DNA sample (representing many millions or even tens of millions of viral genomes (vge) per ml of blood or per ug of total DNA present. We interpret this as evidence of previous persistent infections rather than necessarily currently active systemic infections or shedding as might be detected in the first round cPCR of EHD cases. Therefore, these and similar results should never be interpreted as diagnostic of disease in an asymptomatic healthy elephant, but merely as evidence of a very low-level sporadic persistence following a previous infection that might or might not qualify as periodic reactivation from latency, suggesting further study is needed particularly of possible physical and environmental stressors that may trigger reactivation of EEHV species and subtypes. Severe pathology attributable to EGHV infections including co-morbidity with EHD has also yet to be examined.

## CONCLUSION

We conclude that persistent low-level infections of EEHV2, EEHV3A, EEHV3B, subtypes EEHV3C-EEHV3H, EEHV6, EEHV7A and EEHV7B, and EGHV1B, EGHV2, EGHV3B, EGHV4B and EGHV5B are endemic herpesviruses of *Loxodonta.* Our extensive library of EEHV and EGHV sequences from wild and zoo elephants provides a significant resource to elephant virologists and conservationists worldwide.

## AUTHOR CONTRIBUTIONS

Sample collection, DNA Extraction, cPCR, gel electrophoresis, DNA purification: Pearson

Primer Design: Hayward

Sequencing Analysis: Hayward and Pearson

Original manuscript drafts: Pearson and Hayward

Review & editing: Pearson

## ACKNOWLEDGEMENTS

We thank Lynn W. Enquist, PhD, Princeton University, USA; Glenn F. Rall, PhD, Fox Chase Cancer Center, USA; Jason Holloway, William Rives, DVM, Kenneth Keiffer, DVM, Six Flags Safari Park, USA; Michael Chase, PhD, Kelly Landen, Larry Patterson, DVM, Elephants Without Borders and Government of Botswana; Iain Douglas-Hamilton, PhD, David Daballen, Jerenimo Lepirei, Gilbert Sabinga, Chris Leadismo, Lucy King, PhD, Save The Elephants, and Mathew Mutinda, DVM, Domnic Mjele, DVM, Kenya Wildlife Service Veterinary and Capture Services Department, and Government of Kenya; Michelle Henley, Phd, Elephants Alive, South Africa; Peter Buss, DVM, South African National Parks Veterinary Service and Government of South Africa; Pete Morkel, DVM, Stephanie Bourgeois, PhD, Chimene Nze Nkogue, PhD, Agence National des Parcs Nationaux, CENAREST, and Government of Gabon; Elliott Jacobson, DVM, University of Florida School of Veterinary Medicine, USA; Oliver Ryder, PhD, Christina Reif, San Diego Zoo Wildlife Alliance, USA; Deborah Olson, International Elephant Foundation, USA; Jane Hopper, DVM, Howletts/Port Lympne, England; SY Long, Sarah Heaggens, Johns Hopkins School of Medicine, USA; Albuquerque Zoo, Dickerson Park Zoo, Caldwell Zoo, Cheyenne Mountain Zoo, Cleveland Zoo, Dallas Zoo, Denver Zoo, Fort Worth Zoo, Have Trunk Will Travel, Houston Zoo, Kansas City Zoo, Louisville Zoo, Oklahoma City Zoo, Oregon Zoo, Maryland Zoo in Baltimore; Nashville Zoo, Philadelphia Zoo, Pittsburgh Zoo, St. Louis Zoo and many others with elephants collections, USA; Heidi H.Robbins, Susannah Rouse, the Pearson family, Gentegra LLC, Qiagen Inc., Promega Corporation, Puritan Medical Products.

## Supplemental Information S1: Primers

### SET 1 U38 PAN POL

R1 LGH6711 5’-GTATTTGATTTYGCNAGYYTGTAYCC-3’
R2 LGH6712 5’-TGYAAYGCCGTNTAYGGATTYACCGG-3 L1 LGH6710 5’-ACAAACACGCTGTCRGTRTCYCCRTA-3’
rd1 R1/L1 = 530-bp; rd2 R2/L1 = 250-bp.

### SET 2A U60 PAN TER

R1 LGH6671B 5’-GTTYGTAGTAAANGCCGGATCTAC-3 ‘
R2 LGH3025B 5’-GGTACTATATCTTATCATRTC-3’
R3 LGH-R3alt 5’-GGGGTTTGCCAGGAAGCTAGTG-3’
L2 LGH-L2-2A 5’-GTAATCTCSTATGTGTGYGAGGAG,C-3
L1 5’ LGH7576B 5’-GCNCACGTRAAGAARACACCG,AC-3’ rd1 R1/L1 = 360-bp, rd2 R2/L1 = 290-bp, rd3 R3/L1 = 210-bp.

### SET 4 U71/(gM)

R1 LGH6749 5’-CTATGGGATCCGAACTTTC-3’
R2 LGH6750 5’-CTTTCTAAGGGGGTTTGTTGC-3’
R3 LGH6793 5’-GCCATCGGGATCCGGAAAACGCC-3’
L3 LGH6792 5’-GGGCATCTTCTTCTCCCTCATAGC-3’ L2 LGH6752 5’-CTACATGCCCATGCAGATAGG-3’
rd1 R1/L1 = 750-bp, rd2A R2/L1 and rd2B R1/L2 =730-bp, rd3 R2/L2 = 710-bp, alt rd 2B R1/L3 = 680-bp and alt rd2C R3/L1 = 700-bp;

### SET 5 U51 (vGPCR1) specific for *EEHV2* (not good for *EEHV5*)

R1 LGH7506 5’-GATTGTGAACGCTGTATGTC-3’
R2 LGH7470B 5’-GACAGGTGGTACTGTATGATGTGC-3’ R3 LGH7471 5’-CGGTTACACCGTACCGTGGCTTGC-3’
L3 LGH5201 5’-GCCAGGGTAGATAGAATCAAGGGAA-3’
L2 LGH5200A 5’-CGTGATACGCTTCCAAACATACAGC-3’ L1 LGH4963B 5’-GACTTTCTTCGTGTAGCCCTCGTCTT-3’
rd1 R1/L1 = 910-bp, rd2A R1/L2 = 750-bp, rd3A R1/L3 = 690-bp, and rd2B R2/L1 = 730bp, rd3B R3/L1 = 550-bp or R2/L2 = 570-bp.

### SET 5B U51(vGPCR1) specific for *EEHV1* only (outside pair OK for *EEHV6*)

R1 LGH9323 5’-GTGCTAAACCTTTTCAACGAGACTTC-3’ R2 LGH9351 5’-GCGTTTAACGCMACRCAGTTGGT-3’
L2 LGH9353 5’-CGTAACAGGTTAGCATGAGTATCCTCTTCC-3’ L1 LGH9325 5’-GAGAACGCGTTCTGACTTTCTTCATCA-3’
rd1R1/L1 = 1100-bp; rd2 2A PCR R1/L2 = 990-bp, rd 2B PCR R2/L1 = 1070-bp; rd3AB R2/L2 = 960-bp.

### SET 6 U77 PAN HEL

R1 LGH6649 5’-CCAGTCAACGTATAGCTCGTAG-3
R2 LGH6743 5’-GCAAGGTRGAACGTATCGTCG-3
L3 LGH7990 5’-CACCCACCCARTTGTAGGGAAAGTGC-3’
L2 LGH7885 5’-CTGCGTGTAACATGTGTTC-3’
L1 LGH3198=LGH6742 5’-CACAGMGCGTTGTAGAACC-3’
rd1A R1/L1 = 980-bp, rd2A R1/L2 = 950-bp, rd2B R2/L1 = 685-bp, rd3A R1/L3 = 500-bp, rd3B R2/L2 = 540-bp

### SET 6B U77(HEL) specific for GC-rich *EEHV3/4/7*

R1-7’ LGH7920B 5’-CATGTTSAGGTAGTGCAC**C**GA**G**AGC-3’
R2-7 LGH7921 5’-CTGAACATGTGTTCGGCCAGCAAGG-3’
L2-7 LGH7922 5’-CAGCAGCTAGGAGACGTGGCGAC-3’
L1 LGH3198 5’-CACAGMGCGTTGTAGAACC-3’
rd1 R1-7’/L1 .900-bp, rd2A R1-7’/L2-7 = 500-bp, rd2B R2-7/L1 = 650-bp, rd3AB R2-7/L2-7 = 370-bp.

### SET 7A U60 TER specific for *EEHV3/4*

R1 LGH6707 5’-GTGCTGTAGCGGATCATGTC-3’
R2 LGH6727 5’-GCAACACGAGCACGCAAAGTACGTC-3’ L2 LGH6728 5’-CGGATCATGTCGAACTCCGTG-3’
L0 LGH-LO-7 5’-CGTCGAACACGAGCACGCAAAGTACGTC-3’
rd1 1A R1/L0-7 = 310-bp, rd2A 2A R2/L0-7 = 300-bp, rd2B R1/L2 = 300-bp, rd3A/B R2/L0-7 = 280-bp.

### SET 8 U38 POL specific for *EEHV2*

R1 LGH7440 5’-GACTTCGCCAGCTTGTATCC-3’
R2 LGH7437 5’-GTATCATCAAGCTTATAACC-3’
L2 LGH7450 5’-CTCTACATTTACCGTACACTC-3’
L1 LGH6525 5’-CACATCGATACGGAATCTC-3’
rd1 R1/L1 = 510-bp, rd2A R2/L1 = 490-bp, rd2B R1/L2 = 490-bp, rd3AB R2/L2=470bp.

### SET 8A U38 POL specific for *EEHV2/5*

R1 LGH8423 5’-CATATACGAATGTGCCTCRGAATAYGAGC-3’ R2 LGH8424 5’-GACATGTATCGYGTGTGYATGGATAAGG-3’
L2 LGH8425 5’-CCCGTGTTAGTGGTGACAGTCGTAAC-3’ L1 LGH8426 5’-CCGCCAACCAGGATGTAAGAAGTTG-3’
rd1 R1/L1 = 890-bp, rd2A R1/L2 = 630-bp, rd2B R2/L1 = 555-bp, rd3AB R2/L2 =450-bp.

### SET 9 U38 POL specific for *EEHV3/4*

R1 LGH7400 5’-CAGCATCATCCAGGCCTACAAC-3’
R2 LGH6720 5’-ATCCTGGCGCAGCTGCTGAC-3’
R3 LGH6721 5’-CTCACCTGCAACGCCGTCTA-3’
L1 LGH6719 5’-CGTTGAAGGTGTCGCAGAT-3’
R1’ LGH8750 5’-CTGCTACTACTCCACCCTCGTSCTGG-
L1’ LGH8751 5’-GCCGTGACGGACTCGGCCACGGCCAG-3’
rd1 R1/L1 = 390-bp, rd2 R2/L1 = 270-bp, rd3 R3/L1 = 15-bp, and alt rd2A R1’/L1 = 360-bp, alt rd2B =
R1/L1’ = 330-bp, alt rd3A R1’/L1’ = 300-bp

### SET 12A *EGHV* PAN POL (first step)

R1 LGH6784B 5’-GTGGTKGACTTTGCYAGCCTSTACCC-
R2 LGH6785 5’-CCCMAGYATTATWCAGGCMCA-3’ L1 LGH7489 5’-GTCRGTGTCYCCGTAGAYNAC-3’ rd1 R1/L1-=490
bp, r2 R2/L1 470 bp.

### SET12B POL specific for *EGHV1,2,3,4,5* (second step)

All use rd1 R1/L1, rd2 R2/L1, can add rd3 R2/L2 and R2/L3

**EGHV1**A/B R1-1 5’-CTYTACTGACTCCTGAAGTACTCC-3’
EGHV1A/B R1-2 5’-GYCATCCAGGCCTCAAACCGGGTG-3’
EGHV1A/B L1-2 5’-GTCAAGGCCTGCACCGATGCTGGTG-3’
EGHV1A/B L1-1 5’-CCTAAAGTGGGCTTCGGGATCTCC-3’
**EGHV2** R2-1 5’-CCCTCATAACATCAAAGGAGCTAC-3’
EGHV2 R2-2 5’-CACCCAGAACTAAAGGCGGAGGTG-3’
EGHV2 L2-1 5’-CTTTGATAAGGGATAAGGGATTACAG-3’
**EGHV3**A/B R3-1 5’-CCCAYGAGAAACTACAYATGC-3’
EGHV3A/B R3-2 5’-GCATGGCAATCTAARRCCTGAG-3’
EGHV3A/B L3-1 5’-CTGAATCTGGCATCCGGATCATGTTG-3’
**EGHV4** R4-1 5’-CCCTAATTTCCCACGAGAAACTG-3’ EGHV4 R4-2 5’-CATGCATGGTGACCTAAGGCCCG-3’
EGHV4 L4-1 5’-GACCCTAAATCTTGCATTTGGCTC-3’
EGHV4 NEW L4-2 5’-CTGAGGGCGTCTGGGGTAATTGACTC-3’
EGHV4 NEW L4-3 5’-CACAGGGAAGAACACCAGATGCCACACC-3’
**EGHV5**A/B R5-1 5’-CCCTCATTCCACACAATGAGCTTCATTTGC-3’
EGHV5A/B R5-2 5’-CATTTGCATTCTCATCTTACACCAAACG- EGHV5A/B L5-1 5’- GGGACGACGCAAGGTACGGTATAACCC-3’
Rd2A,2B,3A,3B (multiple alternative combinations)

### SET 15/16A U71/gM specific for *EEHV3/4/7*

R1 LGH6793B 5’-GCCATCGGGATCCGGAAAARACGCC-3’
R2 LGH9209 5’-CCACCGATACTACATGGGAAACATAGCC-3’ (same as LGH9137) R3 LGH9237 5’- CGG GAT TGA AAG GCA AGG ACG GCA ACC C-3’
L4 LGH9211 5’-GGCAACATCACCGTAATGTACGTGGTTTGG-3’
L2 LGH9210 5’CCAGTACGATAAGATCTACCTGGACGA-3’ (same as LGH9138) L1 LGH6792B 5’- GGGCATCTTCTTCACCCTCATAGCCATC-3’
rd1 R1/L1 = 700-bp, rd2 R1/L2 = 670-bp; rd3 R2/L4 PCR = 640-bp.

### SET 16D U71/gM specific for *EEHV3B* and *EEHV7A/B*

R3Bsp LGH9682 5’-GCCTTCGAGGAGCTATCGGATGAAGAGA-3’ L3Bsp LGH9684 5’-GAGGTACAAACCCTTGGTGGTGCTCACA-3’ R3COM LGH9685 5’CTRCTGGACGAC GAAGCCTTCGAGGAGC3’ R7COM LGH9686 5’-CTGTTGGACGACGACGCGTTCGAGGACC-3’
L3+7COM LGH9687 5’-GCGTTGCCTCMGGATTAYTAYCACAAC-3’ Lalt3COM LGH9210 =(9138) 5’-CCAGTACGATAAGATCTACCTGGACGA-3’ rd1 R3/L3+7 =560-bp, rd2 R3B/L3+7 =545-bp, R3B/L3B =510-bp.

### SET 18C SET 18A U73(OBP) specific for GC-rich *EEHV3/4/7*

R1 LGH9279 5’-CGAGCCCCTATGGGGGTCTGGNAAGAC-3’
R1ALT LGH 5’-GGG AGT CGT GAA CCT CGA CAT GAA -3’
R2 LGH9280 5’-GACSGCGGCYATGATCTCGTGGCTCA-3
R6 LGH9382 5’-GTT CCT GGC GTG CGT CGC CTC GTG CC-3’
R7 LGH9394 5’-CTT GCT GCT [C/G]AT GGA CGC CAC C[A/G]T CAA C-3’
L4 LGH9283 5’-GGCACGAGGCGACGCACGCCAGGAACTTG- L0 LGH9288 5’- GACTGGTATACTGACATCATGTCGGGACC-3’
rd1 R1/L0 =940-bp, rd2R1/L4 =360-bp, rd3 R2/L4 = 345-bp
ALT RD1 9382/9288 = 350bp, RD 2 9384/9288 = 345bp

### SET 26 U48.5/U49 (gH-TK)

R1 LGH9985 5’CTGTTCCTCTGTCTCCACGCGCCTCAGCGT-3”
R2 LGH9986 5’-AGCATGGAGTACAACGGCATGTAGCTGA-3’
L2 LGH9987 5’-GTATAGGTGTAGGTAAGACGTCACTGTTC-3’
L1 LGH9988 5’-CTCACCGTTTACCTAGAAGGTTGTATAGGTG-3’
rd1 R1/L1 = 360-bp, rd2A R1/L2 = 300-bp, rd2B R2/L1 = 280-bp, rd3AB R2/L2= 250-bp

### SET 26D new alternative specific for PAN *EEHV3/4/7* GC-branch

R1 LGH11242: 5’-CCTGGCGAAAGGACCAGAGCATGGAG-3’
R2 LGH11243: 5’-GCAGGTACTGGTGCAGGGGAAACAC-3’
L2 LGH11244: 5’-CTGGACCCAGTGGTTTCCCGAAAACG-3’
L1 LGH11245: 5’-CGTCACTGTTCAAATACGCCGCGGATAAC-3’
rd1 R1/L1= 375-bp, rd2A R2/L1 = 320-bp, rd 2B R1/L2 = 305bp, rd 3AB R2/L2 = 250-bp

### SET 27 U81(UDG)

R1 LGH9989 5‘-AGGTAT TGGTTGGCTTTCACGAAGTG-3‘
R2 LGH9990 5‘-CTGGCGGCCGCCAGGGGGGAAGGGTGAGC-3‘
L2 LGH9992 5’-GCCGACGGCCTGGCCTTCTCCACGGGTGACGG-3’
L0 LGH11149 5‘-CGAAGAGTGGGTCGCCTTTCTAGA-3‘
rd 1R1/L0 = 520-bp, rd 2A R1/L2 = 435-bp, rd2B R2/L0= 470-bp, rd R2/L2 = 360-bp;

### SET 28 E54(vOX2-1) (*EEHV1A/1B/6* only) R1

5’ LGH837 5’- ATGCTTCAGAGAAAGTAAGGTAC-3’.
R2alt LGH132 5’-GGTCGTAACGCAAGATGAGCGAG-3’
L1 LGH8472 5’-GTGTTGCCGCCACGATGCTTCTACG-3’
rd1 R1/L1 = 910-bp; rd 2 R2/L1 = 740-bp

### SET 29A U51(vGPCR1) specific for *EEHV3/7*

R1 LGH10784 5’-CGTTCCCGCTGTGGATCTACCAGGC-3’
R2 LGH10785 5’-CAGCGGGATGTTCTACACCATG-3’
R3 LGH10786 5’-GTCACCTTCGATCGCTGGTACTG-3’
L2 LGH10787 5’-GCTCTGTCCCGAGTACAGGTAGCAG-3’ L1 LGH10788 5’-CATGCCGTAGAACGGCCCTTGGG-3’
rd1 R1/L1 = 490bp, rd2A R1/L2 = 310bp,, rd2B R2/L1 = 420, rd3AB R3/L2 =<400bp

### SET 30 E4(vGCNT1) specific for *EEHV4/7*

LGH10789 vGCNT1 EEHV7R1 5’-AGGGCCATCTACGCCCCGCAGAACCT-3’
LGH10790 vGCNT1 EEHV7 R2 5’-TCCTGGAGTCGCGTGCAGGCCGA-3’
LGH10791 vGCNT1 EEHV7 L2 5’-CATGTCCGAGGTGTCGTACTTG-3’
LGH10792 vGCNT1 EEHV7 L1 5’-TCGTAGTAGTGCCACTTGACGAACCTG-3’
rd1 R1/L1 = 660-bp rd2A R2/L1 = 515bp, rd2B R1/L2 = 525bp, rd3AB R2/L2 =415-bp

### SET 35 U14 specific for *EEHV3*

LGHU14-R1 5’-GTCTTCGCAARCCGCGCTGGTCGGC-3’
LGHU14-R2 5’-GTCTCTGGATGTTCTGGGCATCCAG-3’
LGHU14-L2 5’-CCGTGTATGAARTACGTRTGTAAC-3’ LGHU14- L1 5’-CGAGCCGATATCGTCGACGATCTG-3’ rd1 R1/L1 = 920-bp, rd2A R1/L2 =870-bp, rd2B R2/L1 = 860-bp, rd3AB R2/L2 = 810-bp

### SET 36 U42(MTA) specific for *EEHV3A/B* includes across ex1/ex2 boundary

LGHU42-R1 5’-GTAGTCGAAGACAGCGCGTTCTAG-3’
LGHU42-R2 5’-GACAACATGTTCGACACTCCGGTC-3’
LGHU42-L2 5’-GACGTTCGGCAGCTTTCCCGTGTA-3’
LGHU42-L1 5’-GACGAACTTTTTCTCCATGAGCATGG-3’ rd1

R1/L1 = 900bp, rd2A R1/L2 = 860bp, rd2B R1/L2 = 870- bp= rd3AB R2/L2 = 830-bp

### SET 37 U43(PRI) specific for *EEHV3A/B*

LGHU43 R1 5’-CAGGTGCTCTTCGCGACCGAATAC-3’
LGHU43 R2 5’-TGCGGATGCGATGTTGACCACCT-3’
LGHU43 L2 5’-CAGAACGCTATGATCGTTTCGTC- 3’
LGHU43 L1 5’-CTGACGCGGGTAGAGTTTTCGCACG-3’ rd1 R1/L1 = 760bp,

rd2A R1/L2 = 720bp, rd2B R2/L1 = 710bp, rd3AB R2/L2 = 670-bp

### SET 38 U82(gL)-E37(ORF-Oex3) specific for *EEHV3/4/7*

LGH11558 New R1 5’-CGTTCACGGGCGGYGGGTASACGATC-3’
LGH11559 New R2 5’-GGGTASACCGATCTCGGAGACCCG-3’
LGH11560 New L5A 5’-GACGCGTTTATCGTCACRGGGAGGG-3’
LGH11561 New L4 5’-TTYGACACTCCCGCCCTCCTTGACG-3’
LGH11562 New R3 5’-CACYCTYAGTTCCTCSGTRTTATA C-3’
LGH11563 New R4 5’-GAA TAGAYTTAACRCCGGTAGGAAC-3’
LGH11564 NewL3 5’-GTTCCTACCGGYGGTGTTAAATCTATTC-3’
LGH11565 New L2 5’-CAGATGATAACGTCCCTSGACAAGTG-3’
LGH11566 New L1 5’-CAGATYACGTATGTYTTTRATCCG-3

RdA: rd1 R1/L4 =680-bp, rd2A R1/L5 =640-bp, rd2B R2/L4 =66-bp, rd3A R2/L5 =620-bp; Rd2B: rd1 R3/L1=705bp, rd2A R3/L2 =610-bp, rd2B R4/L1 =565-bp, rd3A R3/L3 =535-bp, rd3B R4/L2 =565-bp.

Multiple combinations over 1800-bp.

## REFERENCES

1. Shoshani J. & Tassy P. The Proboscidea: evolution and ptaalaeoecology of elephants and their relatives. Oxford University Press, 1996.

2. IUCN 2025. International Union for Conservation of Nature Red List, icunredlist.org

3. Cerling T. E., Barnette J. E., Chesson L. A., Douglas-Hamilton I., Gobush K. S., Uno K. T., Wasser S. K. & Xu X. Radiocarbon dating of seized ivory confirms rapid decline in African elephant populations and provides insight into illegal trade. Proc Natl Acad Sci U S A 113, 13330–13335, (2016). 10.1073/pnas.1614938113 PMID: 27821744

4. Wittemyer G., Northrup J. M., Blanc J., Douglas-Hamilton I., Omondi P. & Burnham K. P. Illegal killing for ivory drives global decline in African elephants. Proc Natl Acad Sci U S A 111, 13117–13121, (2010.1073/pnas.1403984111 PMID: 25136107

5. Chase M. J., Schlossberg S., Griffin C. R., Bouche P. J., Djene S. W., Elkan P. W., Ferreira S., Grossman F., Kohi E. M., Landen K., Omondi P., Peltier A., Selier S. A. & Sutcliffe R. Continent-wide survey reveals massive decline in African savannah elephants. PeerJ 4, e2354, (2016). 10.7717/peerj.2354 PMID: 27635327

6. Breuer T., Maisels F. & Fishlock V. The consequences of poaching and anthropogenic change for forest elephants. Conserv Biol 30, 1019–1026, (2016). 10.1111/cobi.12679 PMID: 26801000

7. Wasser S. K., Brown L., Mailand C., Mondol S., Clark W., Laurie C. & Weir B. S. CONSERVATION. Genetic assignment of large seizures of elephant ivory reveals Africa’s major poaching hotspots. Science 349, 84–87, (2015). 10.1126/science.aaa2457 PMID: 26089357

8. Maisels F, Strindberg S., Blake S., Wittemyer G., Hart J., Williamson E. A., Aba’a R., Abitsi G., Ambahe R. D., et al. Devastating decline of forest elephants in central Africa. PLoS One 8, e59469, (2013). 10.1371/journal.pone.0059469 PMID: 23469289

9. Poulsen J. R., Koerner S. E., Moore S., Medjibe V. P., Blake S., Clark C. J., Akou M. E., Fay M., Meier A. & Okouyi J. Poaching empties critical Central African wilderness of forest elephants. Current Biology 27, R134–R135(2017). 10.1016/j.cub.2017.01.023 PMID: 28222286

10. Sampson C., McEvoy J., Oo Z. M., Chit A. M., Chan A. N., Tonkyn D., Soe P., Songer M., Williams A. C., Reisinger K., Wittemyer G. & Leimgruber P. New elephant crisis in Asia-Early warning signs from Myanmar. PLoS One 13, e0194113, (2018). 10.1371/journal.pone.0194113 PMID: 29534096

11. Atkins L, Zong JC, Tan J, Mejia A, Heaggans SY, Nofs SA, Stanton JJ, Flanagan JP, Howard L, Latimer E, Stevens MR, Hoffman DS, Hayward GS, Ling PD. Elephant endotheliotropic herpesvirus 5, a newly recognized elephant herpesvirus associated with clinical and subclinical infections in captive Asian elephants (Elephas maximus). J Zoo Wildl Med. 2013 Mar;44(1):136–43. doi: 10.1638/1042-7260-44.1.136. PMID: 23505714; PMCID: PMC3746547.

12. Richman L. K., Montali R. J., Garber R. L., Kennedy M. A., Lehnhardt J., Hildebrandt T., Schmitt D., Hardy D., Alcendor D. J. & Hayward G. S. Novel endotheliotropic herpesviruses fatal for Asian and African elephants. Science 283, 1171–1176, (1999). 10.1126/science.283.5405.1171 PMID: 10024244

13. Stanton JJ, Nofs SA, Zachariah A, Kalaivannan N, Ling PD. Detection of elephant endotheliotropic herpesvirus infection among healthy Asian elephants (Elephas maximus) in South India. J Wildl Dis. 2014 Apr;50(2):279–87. doi: 10.7589/2012-09-236. Epub 2014 Jan 31. PMID: 24484479.

14. Fickel J, Richman LK, Montali R, Schaftenaar W, Göritz F, Hildebrandt TB, Pitra C. A variant of the endotheliotropic herpesvirus in Asian elephants (Elephas maximus) in European zoos. Vet Microbiol. 2001 Sep 20;82(2):103–9. doi: 10.1016/s0378-1135(01)00363-7. PMID: 11423201.

15. Garner M. M., Helmick K., Ochsenreiter J., Richman L. K., Latimer E., Wise A. G., Maes R. K., Kiupel M., Nordhausen R. W., Zong J. C. & Hayward G. S. Clinico-pathologic features of fatal disease attributed to new variants of endotheliotropic herpesviruses in two Asian elephants (Elephas maximus). Vet Pathol 46, 97–104, (2009). 10.1354/vp.46-1-97 PMID: 19112123

16. Ossent P., Guscetti F., Metzler A. E., Lang E. M., Rubel A. & Hauser B. Acute and fatal herpesvirus infection in a young Asian elephant (Elephas maximus). Vet Pathol 27, 131–133, (1990). 10.1177/030098589002700212 PMID: 2161138

17. Zachariah A, Sajesh PK, Santhosh S, Bathrachalam C, Megha M, Pandiyan J, Jishnu M, Kobragade RS, Long SY, Zong JC, Latimer EM, Heaggans SY, Hayward GS. Extended genotypic evaluation and comparison of twenty-two cases of lethal EEHV1 hemorrhagic disease in wild and captive Asian elephants in India. PLoS One. 2018 Aug 22;13(8):e0202438. doi: 10.1371/journal.pone.0202438. PMID: 30133540; PMCID: PMC6105008.

18. Stanton JJ, Nofs SA, Zachariah A, Kalaivannan N, Ling PD. Detection of elephant endotheliotropic herpesvirus infection among healthy Asian elephants (Elephas maximus) in South India. J Wildl Dis. 2014 Apr;50(2):279–87. doi: 10.7589/2012-09-236. Epub 2014 Jan 31. PMID: 24484479.

19. Bauer KL, et al. 2018. Long-term, intermittent, low-level elephant endotheliotropic herpesvirus 1A viremia in a captive Asian elephant calf. Journal of Veterinary Diagnostic Investigation. 30: 917–919.

20. Boonprasert K, Punyapornwithaya V, Tankaew P, Angkawanish T, Sriphiboon S, Titharam C, et al. (2019) Survival analysis of confirmed elephant endotheliotropic herpes virus cases in Thailand from 2006 – 2018. PLoS ONE 14 (7): e0219288. 10.1371/journal.pone.0219288

21. Bouchard B., Xaymountry B., Thongtip N., Lertwatcharasarakul P. & Wajjwalku W. First reported case of elephant endotheliotropic herpes virus infection in Laos. J Zoo Wildl Med 45, 704–707, (2014). 10.1638/2013-0264R1.1 PMID: 25314848

22. Dastjerdi A., Seilern-Moy K., Darpel K., Steinbach F. & Molenaar F. Surviving and fatal Elephant Endotheliotropic Herpesvirus-1A infections in juvenile Asian elephants—lessons learned and recommendations on anti-herpesviral therapy. BMC veterinary research 12, 178, (2016). 10.1186/s12917-016-0806-5 PMID: 27567895

23. Fuery A., Tan J., Peng R., Flanagan J. P., Tocidlowski M. E., Howard L. L. & Ling P. D. Clinical Infection of Two Captive Asian Elephants (Elephas Maximus) with Elephant Endotheliotropic Herpesvirus 1b. J Zoo Wildl Med 47, 319–324, (2016). 10.1638/2015-0074.1 PMID: 27010294

24. Lee MH, Nathan SKSS, Benedict L, Nagalingam P, Latimer E, Hughes T, Ramirez D, Sukor JRA. The first reported cases of elephant endotheliotropic herpesvirus infectious haemorrhagic disease in Malaysia: case report. Virol J. 2021 Nov 24;18(1):231. doi: 10.1186/s12985-02101694-x. PMID: 34819101; PMCID: PMC8611640

25. Oo Z. M., Aung Y. H., Aung T. T., San N., Tun Z. M., Hayward G. S. & Zachariah A. Elephant Endotheliotropic Herpesvirus Hemorrhagic Disease in Asian Elephant Calves in Logging Camps, Myanmar. Emerg Infect Dis 26, 63–69, (2020). 10.3201/eid2601.190159 PMID: 31855135

26. Pavulraj S., Eschke K., Prahl A., Flügger M., Trimpert J., van den Doel P. B., Andreotti S., Kaessmeyer S., Osterrieder N. & Azab W. Fatal Elephant Endotheliotropic Herpesvirus Infection of Two Young Asian Elephants. Microorganisms 7, (2019). 10.3390/microorganisms7100396 PMID: 31561506

27. Reid C. E., Hildebrandt T. B., Marx N., Hunt M., Thy N., Reynes J. M., Schaftenaar W. & Fickel J. Endotheliotropic elephant herpes virus (EEHV) infection. The first PCR-confirmed fatal case in Asia. Vet Q 28, 61–64, (2006). 10.1080/01652176.2006.9695209 PMID: 16841568

28. Sripiboon S, Tankaew P, Lungka G, Thitaram C. The occurrence of elephant endotheliotropic herpesvirus in captive Asian elephants (Elephas maximus): first case of EEHV4 in Asia. J Zoo Wildl Med. 2013 Mar;44(1):100–4. doi: 10.1638/1042-7260-44.1.100. PMID: 23505709.

29. Stremme C, Priadi A, Hayward GS, Zachariah A. IDENTIFICATION OF TWO LETHAL CASES OF ELEPHANT ENDOTHELIOTROPIC HERPESVIRUS HEMORRHAGIC DISEASE IN SUMATRAN ELEPHANT CALVES IN INDONESIA. J Zoo Wildl Med. 2021 Jan;51(4):985–993. doi: 10.1638/2020-0003. PMID: 33480579.

30. Wilkie G. S., Davison A. J., Kerr K., Stidworthy M. F., Redrobe S., Steinbach F., Dastjerdi A. & Denk D. First fatality associated with elephant endotheliotropic herpesvirus 5 in an Asian elephant: pathological findings and complete viral genome sequence. Sci Rep 4, 6299, (2014). 10.1038/srep06299 PMID 25199796

31. Yun Y, Sripiboon S, Pringproa K, Chuammitri P, Punyapornwithaya V, Boonprasert K, Tankaew P, Angkawanish T, Namwongprom K, Arjkumpa O, Brown JL, Thitaram C. Clinical characteristics of elephant endotheliotropic herpesvirus (EEHV) cases in Asian elephants (*Elephas maximus*) in Thailand during 2006-2019. Vet Q. 2021 Dec;41(1):268–279. doi: 10.1080/01652176.2021.1980633. PMID: 34511026; PMCID: PMC847

32. Zachariah A., Zong J. C., Long S. Y., Latimer E. M., Heaggans S. Y., RiHayward G. S. Fatal herpesvirus hemorrhagic disease in wild and orphan Asian elephants in southern India. J Wildl Dis 49, 381–393, (2013). 10.7589/2012-07-193 PMID: 23568914

33. Ackermann M, Hatt JM, Schetle N, Steinmetz H. Identification of shedders of elephant endotheliotropic herpesviruses among Asian elephants (Elephas maximus) in Switzerland. PLoS One. 2017 May 3;12(5):e0176891. doi: 10.1371/journal.pone.0176891. PMID: 28467495; PMCID: PMC5415103.

34. Perrin KL, Kristensen AT, Bertelsen MF, Denk D. Retrospective review of 27 European cases of fatal elephant endotheliotropic herpesvirus-haemorrhagic disease reveals evidence of disseminated intravascular coagulation. Sci Rep. 2021 Jul 8;11(1):14173. doi: 10.1038/s41598-021-93478-0. PMID: 34238966; PMCID: PMC8266883.

35. Bronson E., McClure M., Sohl J., Wiedner E., Cox S., Latimer E. M., Pearson V. R., Hayward G. S., Fuery A. & Ling P. D. Epidemiologic Evaluation of Elephant Endotheliotropic Herpesvirus 3B Infection in an African Elephant (Loxodonta Africana). J Zoo Wildlife Med 48, 335–343, (2017). 10.1638/2016-0063R.1 PMID: 28749266

36. Melissa A Fayette ^1^, Emily E Brenner ^2^, Michael M Garner ^3^,Michelle R Bowman ^2^, Erin Latimer ^4^, Jeffry S Proudfoot ^2^ ACUTE HEMORRHAGIC DISEASE DUE TO ELEPHANT ENDOTHELIOTROPIC HERPESVIRUS 3A INFECTION IN FIVE AFRICAN ELEPHANTS ( *LOXODONTA AFRICANA*) AT ONE NORTH AMERICAN ZOOLOGICAL INSTITUTION. J Zoo Wildl Med 2021 Apr;52(1):357–365. doi: 10.1638/2020-0126.

37. Fayette MA, Minich DJ, Sylvester H, Latimer E. FIRST DETECTION OF CLINICAL DISEASE DUE TO ELEPHANT ENDOTHELIOTROPIC HERPESVIRUS 7A IN TWO AFRICAN ELEPHANTS (*LOXODONTA AFRICANA*) IN HUMAN CARE. J Zoo Wildl Med. 2024 Mar;55(1):290–294. doi: 10.1638/2023-0034. PMID: 38453514.

38. Knüppel, S.; Balfanz, F.; Riedel, C.; Strauss, V.; Hoornweg, T.E.; Dimmel, K.; Walk, K.; Kübber-Heiss, A.; Posautz, A.; Voracek, T.;, et al. Severe Elephant Endotheliotropic Herpesvirus 6 Associated Disease in Two African Elephants Under Human Care in Austria. Animals 2025, 15, 1482. 10.3390/ani15101482.

39. Willis TJ, Burgdorf-Moisuk A, Connolly M, Raines J, Ratliff C, Gyimesi ZS, Lipanovich E, Dumonceaux GA, Perrin KL, Howard LL, Kinney ME, Suedmeyer K, Latimer E, Kim TL, Watts JR, Tan J, Ling P. Elephant endotheliotropic herpesvirus 2 infection in 5 African elephants (Loxodonta africana) at multiple North American zoological institutions. Am J Vet Res. 2025 Jul 2;86(9):ajvr.25.02.0060. doi: 10.2460/ajvr.25.02.0060. PMID: 40602616

40. Kongmakee, P., Suttiyaporn, S., Changpetc, W., Kongkham, W., Mongkolphan, C., Tonchiangsai, K. & et al. in Proceedings Tenth International Elephant Endotheliotropic Herpesvirus (EEHV) Workshop 2015.

41. Pursell T, Spencer Clinton JL, Tan J, Peng R, Qin X, Doddapaneni H, Menon V, Momin Z, Kottapalli K, Howard L, Latimer E, Heaggans S, Hayward GS, Ling PD. Primary Infection May Be an Underlying Factor Contributing to Lethal Hemorrhagic Disease Caused by Elephant Endotheliotropic Herpesvirus 3 in African Elephants (*Loxodonta africana*). Microbiol Spectr. 2021 Oct 31;9(2):e0098321. doi: 10.1128/Spectrum.00983-21. Epub 2021 Oct 20. PMID: 34668724; PMCID: PMC8528115

42. Kerr TJ, van Heerden J, Goosen WJ, Kleynhans L, Buss PE, Latimer E, Miller MA. DETECTION OF ELEPHANT ENDOTHELIOTROPIC HERPESVIRUS (EEHV) IN FREE-RANGING AFRICAN ELEPHANTS (LOXODONTA AFRICANA) IN THE KRUGER NATIONAL PARK, SOUTH AFRICA. J Wildl Dis. 2023 Jan 1;59(1):128–137. doi: 10.7589/JWD-D-22-00015. PMID: 36584337.

43. Howard LL, A Primer on Elephant Endothelitropic Herpesvirus (EEHV), 2025, www.eehvinfo.org

44. Zong J. C., Heaggans S. Y., Long S. Y., Latimer E. M., Nofs S. A., Bronson E., Casares M., Fouraker M. D., Pearson V. R., Richman L. K. & Hayward G. S. Detection of Quiescent Infections with Multiple Elephant Endotheliotropic Herpesviruses (EEHVs), Including EEHV2, EEHV3, EEHV6, and EEHV7, within Lymphoid Lung Nodules or Lung and Spleen Tissue Samples from Five Asymptomatic Adult African Elephants. J Virol 90, 3028–3043, (2015). 10.1128/JVI.02936-15 PMID: 26719245

45. Jacobson E. R., Sundberg J. P., Gaskin J. M., Kollias G. V. & O’Banion M. K. Cutaneous papillomas associated with a herpesvirus-like infection in a herd of captive African elephants. J Am Vet Med Assoc 189, 1075–1078 (1986). PMID: 2851.

46. McCully, R.M., Basson, P.A., Pienaar, J.G., Erasmus, B.J., Young, E., 1971. Herpes nodules in the lung of the African elephant (Loxodonta africana (Blumebach, 1792)). Onderstepoort J Vet Res 38, 225–235.

47. Pearson, VR, Finding EEHVs in Wild Kenyan Elephants, Journal of Elephant Managers Association, 2012, Volume 23, Number 2 pp.6–8. www.elephantmanagers.org

48. Pearson, VR, International Elephant and Rhino Conservation and Research Symposium, 2013, Pittsburgh, PA, USA, pp. 592– 741

49. Pearson, VR, Ninth International Workshop for Elephant Endothelioltropic Herpesviruses, 2013, (Houston, TX, USA, 2013).

50. Pearson VR, Zong JC, Heaggans SY, Long SY & Hayward GS. Genetic Characterization of EEHV and Gammaherpesviruses in Lung or Skin Nodules and Saliva from Zoo and Wild African Elephants. 2016, IEF/IRF Research Symposium, Singapore.

51 Landolfi JA, Howard L, Ling P (2025) Tissue and cellular tropism of elephant endotheliotropic herpesvirus (EEHV)1A in hemorrhagic disease. PLoS One 20(9): e0330631. 10.1371/journal.pone.0330631

52. Stanton JJ, Zong JC, Latimer E, Tan J, Herron A, Hayward GS, Ling PD. Detection of pathogenic elephant endotheliotropic herpesvirus in routine trunk washes from healthy adult Asian elephants (Elephas maximus) by use of a real-time quantitative polymerase chain reaction assay. Am J Vet Res. 2010 Aug;71(8):925–33. doi: 10.2460/ajvr.71.8.925. PMID: 20673092; PMCID: PMC3725808.

53. Jeffrey A, et al. 2020. Noninvasive sampling for detection of elephant endotheliotropic herpesvirus and genomic DNA in Asian (Elephas maximus) and African (*Loxodonta africana*) elephants. Journal of zoo and wildlife medicine. 51:433–437.

54. Common SM, et al. 2021. Developing a non-invasive method of detecting elephant endotheliotropic herpesvirus infections using faecal samples. Veterinary Record. e833. 10.1002/vetr.833 Gentry PA, et al. 1996. Blood coagulation profile of the Asian elephant (*Elephas maximus*). Zoo Biology 15: 413–423.

55. Sylvester H, Raines J, Burgdorf-Moisuk A, Connolly M, Wilson S, Ripple L, Rivera S, McCain S, Latimer E. SELECTED INSTANCES OF ELEPHANT ENDOTHELIOTROPIC HERPESVIRUS SHEDDING IN TRUNK SECRETIONS BY AFRICAN ELEPHANTS (*LOXODONTA AFRICANA*) IN COMPARISON TO SHEDDING BY ASIAN ELEPHANTS (*ELEPHAS MAXIMUS*). J Zoo Wildl Med. 2024 Mar;55(1):182–194. doi: 10.1638/2022-0046. PMID: 38453501.

56. Grenus BG, Latimer E, Cullinane A, Lyons P, Creighton G, Nutter FB. EVALUATION OF THE EFFICACY OF TWO DIFFERENT SAMPLING SITES FOR THE DETECTION OF ELEPHANT ENDOTHELIOTROPIC HERPESVIRUS (EEHV) IN THREE ASIAN ELEPHANTS (*ELEPHAS MAXIMUS*) IN IRELAND. J Zoo Wildl Med. 2020 Jun;51(2):303–307. doi: 10.1638/2018-0193. PMID: 32549559.

57. Yang N, Bao M, Zhu B, Shen Q, Guo X, Li W, Tang R, Zhu D, Tang Y, Phalen DN, Zhang L. Elephant Endotheliotropic Herpesvirus 1, 4 and 5 in China: Occurrence in Multiple Sample Types and Implications for Wild and Captive Population Surveillance. Viruses. 2022 Feb 17;14(2):411. doi: 10.3390/v14020411. PMID: 35216004; PMCID: PMC8875873.

58. Oliveira Neto NF, Caixeta RAV, Zerbinati RM, Zarpellon AC, Caetano MW, Pallos D, Junges R, Costa ALF, Aitken-Saavedra J, Giannecchini S, Braz-Silva PH. The Emergence of Saliva as a Diagnostic and Prognostic Tool for Viral Infections. Viruses. 2024 Nov 11;16(11):1759. doi: 10.3390/v16111759. PMID: 39599873; PMCID: PMC11599014.

59. Robbins HH, Elephant Endotheliotropic Herpesviruses in African Elephants and Detection by Saliva Sampling, Undergraduate Thesis 2013, Princeton University, Princeton, NJ

60. Pearson VR, Bosse JB, Koyuncu OO, Scherer J, Toruno C, Robinson R, et al. (2021) Identification of African Elephant Polyomavirus in wild elephants and the creation of a vector expressing its viral tumor antigens to transform elephant primary cells. PLoS ONE 16(2): e0244334. 10.1371/journal.pone.024433

61. Ling PD, Long SY, Zong J, Heaggans SY, Qin X, Hayward GS.2016. Comparison of the Gene Coding Contents and Other Unusual Features of the GC-Rich and AT-Rich Branch Probosciviruses. mSphere1:10.1128/msphere.00091-16.10.1128/msphere.00091-16

62. Ling PD, Long SY, Fuery A, Peng R, Heaggans SY, Qin X, Worley KC, Dugan S, Hayward GS.2016.Complete Genome Sequence of Elephant Endotheliotropic Herpesvirus 4, the First Example of a GC-Rich Branch Proboscivirus. mSphere1:10.1128/msphere.00081-15.10.1128/msphere.00081-15

63. Zong J. C., Latimer E. M., Long S. Y., Richman L. K., Heaggans S. Y. & Hayward G. S. Comparative genome analysis of four elephant endotheliotropic herpesviruses, EEHV3, EEHV4, EEHV5, and EEHV6, from cases of hemorrhagic disease or viremia. J Virol 88, 13547–13569, (2014). 10.1128/JVI.01675-14 PMID: 25231309

64. Tan, J., Ling, P.D., Worley, K., Proudfoot, J., Bowman, M., Qin, X., Latimer, E.M., Holder, K., Fayette, M., Nodolf, S., Heaggans, S.Y., Zong, J.-C., Pearson, V.R. and Hayward, G.S. Complete Genome Assembly and Annotation of EEHV3A the First Example of a GC-Branch African Elephant Endotheliotrophic Herpesvirus Associated with Lethal Hemorrhagic Disease, manuscript in preparation.

65. Richman LK, Zong JC, Latimer EM, Lock J, Fleischer RC, Heaggans SY, Hayward GS. J Virol. 2014 Dec;88(23):13523–46. doi: 10.1128/JVI.01673-14. Elephant endotheliotropic herpesviruses EEHV1A, EEHV1B, and EEHV2 from cases of hemorrhagic disease are highly diverged from other mammalian herpesviruses and may form a new subfamily.

66. Ehlers B, Dural G, Yasmum N, Lembo T, de Thoisy B, Ryser-Degiorgis MP, Ulrich RG, McGeoch DJ. Novel mammalian herpesviruses and lineages within the Gammaherpesvirinae: cospeciation and interspecies transfer. J Virol. 2008 Apr;82(7):3509–16. doi: 10.1128/JVI.0264607. Epub 2008 Jan 23. PMID: 18216123; PMCID: PMC2268488.

67. Latimer E., Zong J. C., Heaggans S. Y., Richman L. K. & Hayward G. S. Detection and evaluation of novel herpesviruses in routine and pathological samples from Asian and African elephants: identification of two new probosciviruses (EEHV5 and EEHV6) and two new gammaherpesviruses (EGHV3B and EGHV5). Vet Microbiol 147, 28–41, (2011). 10.1016/j.vetmic.2010.05.042 PMID: 20579821

68. Masters N. J., Stidworthy M. F., Everest D. J., Dastjerdi A. & Baulmer S. Detection of EGHV-5 in a self-limiting papilloma-like lesion in the trunk of an Asian elephant (Elephas maximus). Vet Rec 169, 209, (2011).

69. Wellehan J. F., Johnson A. J., Childress A. L., Harr K. E. & Isaza R. Six novel gammaherpesviruses of Afrotheria provide insight into the early divergence of the Gammaherpesvirinae. Vet Microbiol 127, 249–257, (2008). 10.1016/j.vetmic.2007.08.024 PMID: 17884307

70. Hayward G. S. Conservation: clarifying the risk from herpesvirus to captive Asian elephants. Vet Rec 170, 202–203, (2012). 10.1136/vr.e1212 PMID: 22368209

71. Long S. Y., Latimer E. M. & Hayward G. S. Review of Elephant Endotheliotropic Herpesviruses and Acute Hemorrhagic Disease. ILAR J 56, 283–296, (2016). 10.1093/ilar/ilv041 PMID: 26912715, 23568914

72. Zong J-C^1^, Pearson VR^2^, Zachariah A^3^, Krushnankutty S^4^, Qin Z^5^, Menon VK^5^, Latimer EM^6^, Heaggans SY^1^, Atkins L^7^, Ling PD^7^ and Hayward GS^1^. UPDATE ON THE STATUS OF EEHV GENOMICS AND ANALYSIS OF SUBTYPES AND STRAIN VARIABILITY: THE “EEHV UNIVERSE” PROJECT., manuscript in preparation.

